# VIA1 is a conserved regulator of thylakoid membrane integrity that acts through VIPP1

**DOI:** 10.64898/2026.03.26.714412

**Authors:** Pamela Vetrano, Kelsey Krall, Laura Martinez, Eleonora Traverso, Tomas Morosinotto, Nicholas A. T. Irwin, Yuval Mazor, Silvia Ramundo

## Abstract

Thylakoid membranes are indispensable for oxygenic photosynthesis, yet the mechanisms that protect these membranes from photooxidative damage remain poorly understood. By screening previously uncharacterized proteins induced during the chloroplast unfolded protein response, we identify VIA1 as an essential factor for preserving thylakoid integrity under high light in the model green alga *Chlamydomonas reinhardtii*. Loss of VIA1 causes hypersensitivity to photo-oxidative stress and rapid thylakoid swelling. VIA1 localizes to thylakoid membranes and directly binds Vesicle-Inducing Protein in Plastids 1 (VIPP1), an ESCRT-III–like protein essential for thylakoid biogenesis and remodeling. Structure-guided mutagenesis shows that this interaction is required for VIA1 function and is mediated by a winged-helix domain interface reminiscent of ESCRT-II/ESCRT-III binding mode. VIA1 orthologs from cyanobacteria and land plants rescue the Chlamydomonas *via1* mutant phenotype, and disruption of VIA1 in *Synechocystis sp*. PCC 6803 impairs growth, especially under light stress. Together, these findings establish VIA1 as an evolutionarily conserved protein that contributes to thylakoid membrane homeostasis via its interaction with VIPP1.

**Significance Statement:** From cyanobacteria to land plants, all organisms performing oxygenic photosynthesis rely on thylakoid membranes to capture light and and produce oxygen. Yet these membranes are highly susceptible to environmental stress, particularly excess light, which causes oxidative damage to membrane lipids and proteins. How thylakoid integrity is maintained under these conditions remains a key open question. Here we identify VIA1 as a conserved factor required for maintaining thylakoid membrane structure under high light. VIA1 interacts with VIPP1, an ESCRT-III-like protein essential for thylakoid biogenesis, through a functionally indispensable interface reminiscent of ESCRT-II/ESCRT-III binding mode. The conservation of the VIA1–VIPP1 module across photosynthetic prokaryotes and eukaryotes suggests it arose early in the evolution of oxygenic photosynthesis and has been maintained ever since.

## Introduction

Photosynthesis, essential for much of life on Earth, is dependent on the extensive network of highly dynamic and specialized membranes known as thylakoids. This membrane system predates the origin of chloroplasts, as it is also present in cyanobacteria. While its composition has remained highly conserved through evolution^1^, the spatial organization of thylakoids has undergone significant change over time^2,3^, suggesting that their biogenesis has also evolved. The complexity of this process—which involves the coordinated synthesis, assembly, and integration of lipids, proteins, and pigments—and its susceptibility to external stress stimuli, has impeded efforts to elucidate the underlying mechanisms. In fact, over the past twenty years, only a few of the molecular components directly involved in the biogenesis and maintenance of thylakoid membranes have been discovered^4–10^.

One of the most extensively investigated of these factors is Vesicle Inducing Protein in Plastid 1 (VIPP1), an essential protein found in nearly all oxygenic photosynthetic organisms^11^. Several studies have pointed to VIPP1’s involvement in thylakoid biogenesis^4,11–13^ as well as in chloroplast envelope and thylakoid membrane maintenance^14–17^. Most notably, the deletion of the *VIPP1* gene in the land plant *Arabidopsis thaliana* and the cyanobacterium *Synechocystis sp.* PCC6803 was shown to induce severe abnormalities in their thylakoid membrane architecture^4,11,15^. Another study in the unicellular green alga *Chlamydomonas reinhardtii* demonstrated that downregulation of VIPP1 via RNA interference causes light-induced thylakoid swelling^14^. A similar phenotype was also observed in Arabidopsis chloroplasts upon knockdown of VIPP1^15^ and in Synechocystis cells expressing a mutant variant of VIPP1^18^.

Recent studies on cyanobacterial VIPP1 have revealed that this protein shares an evolutionary origin and structural features with the Endosomal Sorting Complex Required for Transport (ESCRT)-III components of eukaryotic cells^18,19^. The ESCRT machinery mediates membrane remodeling events in a wide range of cellular contexts and was long thought to be restricted to eukaryotes. However, the discovery of ESCRT-III–like proteins in archaea, cyanobacteria, and chloroplasts has uncovered an ancient, conserved membrane-remodeling module that likely predates the emergence of eukaryotic cells^18–22^. Despite these advances, how VIPP1 is controlled *in vivo*—and whether it acts with dedicated partners to preserve thylakoids, particularly under stress—remains poorly understood.

To address this question, we leveraged the chloroplast unfolded protein response (cpUPR) to identify VIPP1-interacting proteins that contribute to thylakoid membrane integrity and photoprotection. In Chlamydomonas, cpUPR activation—triggered by artificial repression of an essential chloroplast protease, or by physiological photoxidative stress—drives a nuclear gene expression program that requires the cytosolic Ser/Thr kinase MARS1^23^. During this response, VIPP1 is induced alongside protein chaperones, proteases, and multiple poorly annotated genes, suggesting a coordinated effort to stabilize the highly protein-rich thylakoid membranes when chloroplast proteostasis is challenged. This observation prompted us to examine uncharacterized cpUPR-induced factors for their potential roles in membrane homeostasis. Among these, we focused on CPLD50, which we renamed VIPP1-Associated Protein 1 (VIA1)^24,25^. VIA1 is conserved from cyanobacteria to land plants and contains two winged-helix (WH) domains, a fold typically found in ESCRT-II proteins and known to mediate interactions with ESCRT-III proteins^26^.

Here, we show that *via1* hypomorphic mutants are hypersensitive to photo-oxidative stress and exhibit thylakoid membrane perturbations specifically under high light exposure, reminiscent of phenotypes reported for VIPP1 knockdown in Chlamydomonas. Mechanistically, we demonstrate that VIA1 binds VIPP1 directly via its WH domains, and that this interaction is required for VIA1 function. Moreover, the VIA1–VIPP1 module is functionally conserved, as VIA1 orthologs from Synechocystis and Arabidopsis rescue a Chlamydomonas *via1* mutant. Thus, VIA1 is a conserved regulator of VIPP1-dependent membrane remodeling.

## Results

### VIA1 (CPLD50) is a highly conserved protein upregulated upon photooxidative stress in a MARS1-dependent manner

We reasoned that proteins acting alongside VIPP1 to preserve thylakoid structural integrity should be conserved across photosynthetic lineages, chloroplast-localized, and capable of associating with membranes. We therefore prioritized candidates with these characteristics among the MARS1-dependent genes activated during the cpUPR. CPLD50 (hereafter VIA1) emerged as a strong hit for three reasons: (i) it belongs to the GreenCut^27^, a group of nuclear-encoded proteins restricted to green-lineage organisms, consistent with a photosynthesis- or chloroplast-associated function; (ii) it has been previously detected in proteomic surveys aimed at defining chloroplast-localized proteins and is also predicted to be chloroplast-targeted (TargetP2.0^28^/WoLF PSORT^29^/DeepLoc^30^); and (iii) it is predicted to encode three transmembrane helices, suggesting membrane association.

To confirm that VIA1 expression is stress-responsive, we performed RT–qPCR analysis in wild type (WT) and *mars1* knockout (*mars1*) cultures exposed to high light (HL) and monitored the expression of VIA1 alongside canonical cpUPR marker genes (SNOAL2, VIPP2, HSP22E). In WT cells, all tested genes—including VIA1—were strongly induced by photo-oxidative stress, whereas this induction was abolished in the *mars1* mutant. These results confirmed that VIA1 is activated by HL in a MARS1-dependent manner (Fig. S1).

Next, to validate and extend the Green Cut assignment, we reconstructed a VIA1 phylogeny and assessed its distribution across eukaryotic and prokaryotic lineages (Fig. 1A–B). VIA1 is highly conserved across most photosynthetic eukaryotes, including species bearing primary and secondary plastids, but has been frequently lost in photosynthetic dinoflagellates. This distribution, together with the presence of homologs in cyanobacteria, supports a plastid-associated origin for VIA1 dating back to the primary cyanobacterial endosymbiosis in the Viridiplantae.

**Figure 1.**
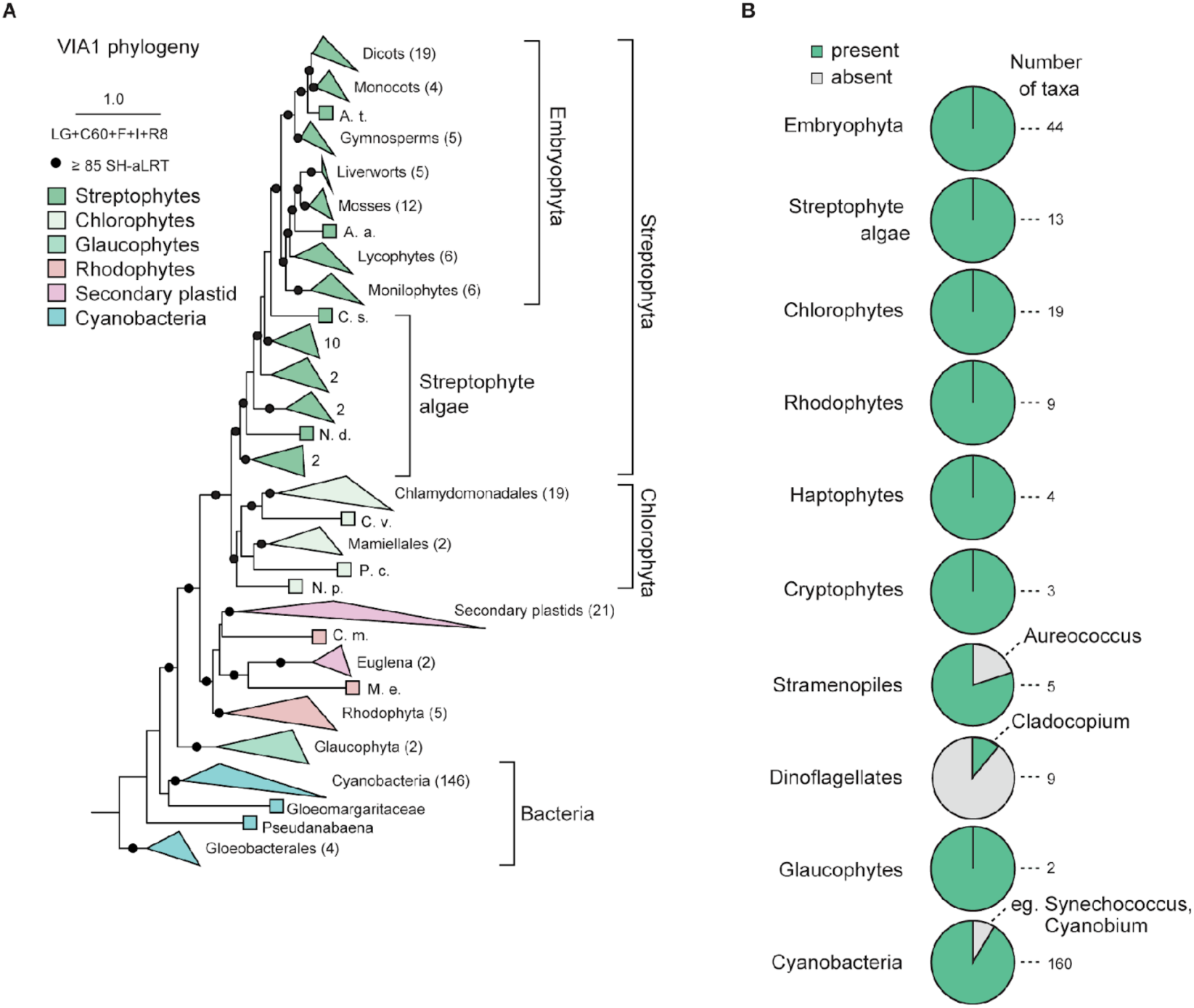
VIA1 is an evolutionarily conserved protein across photosynthetic lineages. **(A)** Maximum-likelihood phylogeny of VIA1. Taxonomic groups have been coloured as noted, and triangles reflect collapsed clades with the total number of taxa denoted in parentheses. Scale bar reflects the average number of substitutions per site. SH-aLRT, Shimodaira-Hasegawa approximate likelihood ratio test. **(B)** Presence of VIA1 across key photosynthetic lineages. The full phylogeny and species distribution are available at https://itol.embl.de/shared/1WjO4fmjYiy3z

### Loss of VIA1 leads to thylakoid structural alterations under high light and renders Chlamydomonas hypersensitive to photooxidative stress

To address the functional role of VIA1, we obtained two independent insertional *via1* mutant strains of *Chlamydomonas reinhardtii*, hereafter referred to as *via1-1* and *via1-2*, from the Chlamydomonas Library Project (CLiP)^31^. The insertion sites within the VIA1 locus were confirmed by PCR (Fig. S2 B–D) and backcross analyses (Fig. S2E-F). Moreover, RT-qPCR (Fig. 2A–B) suggested that VIA1 expression is reduced in both *via1-1* and *via1-2*.

**Figure 2.**
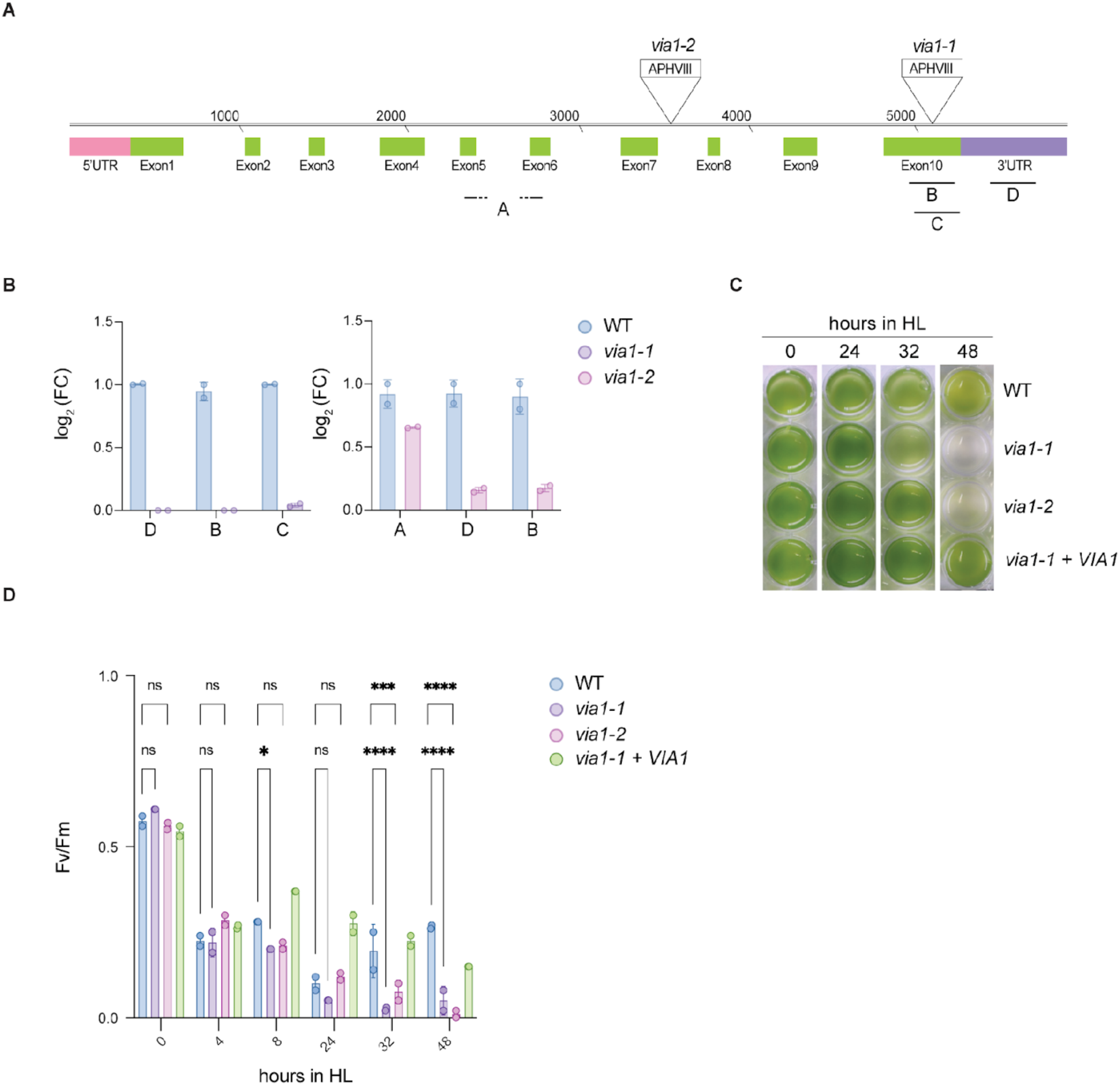
VIA1 is required for high light acclimation. **(A)** VIA1 gene model showing exons, 5′ and 3′ untranslated regions (UTRs). The position of the mutagenic cassette carrying the paromomycin resistance gene (*APHVIII*) in *via1-1* and *via1-2* is indicated. Letters (A–D) denote the amplicon regions used for RT–qPCR in (B). **(B)** VIA1 transcript levels in *via1-1* and *via1-2*, quantified by RT–qPCR using the regions indicated in (A). *GBLP* served as the reference gene for normalization. **(C)** Representative images of liquid TAP cultures of WT, *via1-1*, *via1-2*, and *via1-1* + VIA1 at various time points during a 48-h HL treatment. **(D)** Photosystem II quantum yield (Fv/Fm) measured over 48 h of HL exposure in WT, *via1-1*, *via1-2*, and *via1-1* + VIA1 strains. The significance was assessed via Two-Way ANOVA and Tukey’s multiple comparisons analysis.

When grown under standard light conditions, *via1-1* and *via1-2* mutants were indistinguishable from WT. This was also supported by measurements of key photosynthetic parameters, including the quantum yield of photosystem I (Y_I_), the maximum quantum efficiency of photosystem II (Fv/Fm), the plastoquinone redox state (1-qL), and non-photochemical quenching (NPQ) (Fig. S3 A to D), as well as by growth assays on TAP agar (Fig. S3 E–F). In contrast, when exposed to HL, both mutants were significantly more sensitive to photooxidative stress than WT. In particular, chlorophyll content was significantly reduced in mutants compared to WT after 2 days, and photobleaching occurred earlier in the mutants (Fig. 2C; Fig. S4A–B). Consistently, Fv/Fm declined significantly in both mutants after approximately 32 hours of HL exposure (Fig. 2D). These data rule out a primary photosynthetic defect in either mutant while demonstrating that VIA1 is essential for maintaining photosynthetic function under photooxidative stress.

To determine the underlying cause for the HL sensitivity in *via* mutants, we examined chloroplast ultrastructure by transmission electron microscopy (TEM). Thylakoids in *via1* cells became severely swollen as early as 4 h after the HL onset (Fig. 3B), whereas they remained largely unaffected in WT until 6 h (Fig. 3A and Fig. S4C). This phenotype resembled the HL-induced thylakoid defects reported for VIPP1 knockdown^14^, albeit with delayed onset. To assess whether photosynthetic protein levels were also affected, we examined steady-state levels of core subunits of photosystem I, photosystem II, cytochrome b₆f, and ATP synthase by immunoblotting at 2-hour intervals over an 8-hour HL time course. This analysis revealed comparable steady-state levels of core subunits of photosystem I, photosystem II, cytochrome *b*₆*f*, and ATP synthase in WT and *via1* cells throughout the time course (Fig. S4D). Thus, these results suggest that VIA1’s primary function is to maintain thylakoid membrane integrity under photooxidative stress, whereas the decline in photosynthetic activity observed at later time points is likely a secondary consequence.

**Figure 3.**
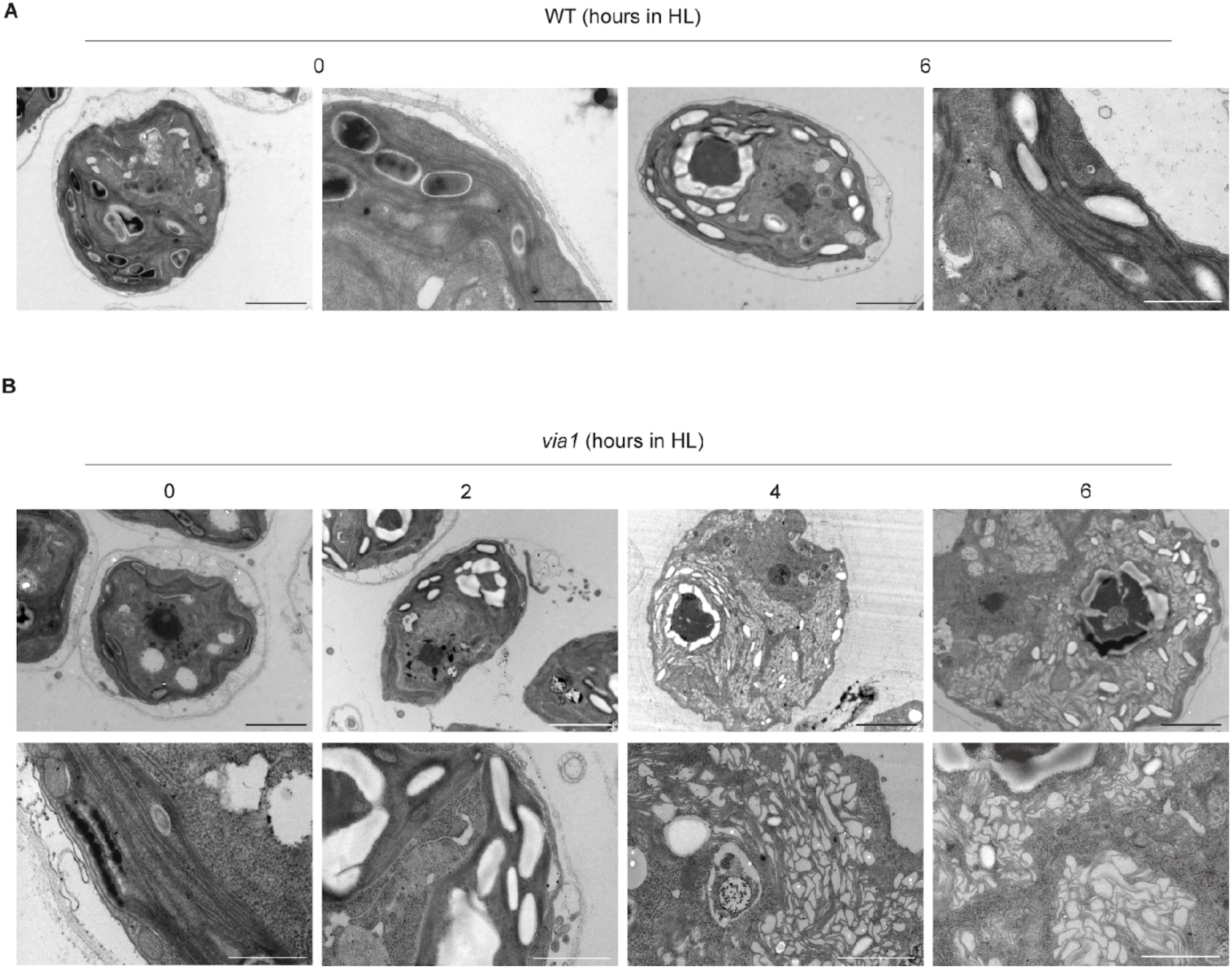
VIA1 is required for thylakoid membrane stability. **(A)** Representative transmission electron microscopy (TEM) images of WT cells captured at 0 and 6 h of HL exposure (∼1000 µmol photons m^−2^ s^−1^). Scale bars correspond to 2 µm for whole-cell images and 1 µm for zoomed-in images. **(B)** Representative TEM images of *via1-2* cells at 0-, 2-, 4-, and 6-hours of HL exposure (∼1000 µmol photons m^−2^ s^−1^). Scale bars correspond to 2 µm for the first row, 500 nm for the first image in the lower row, and 1 µm for the remaining images.

### VIA1 is part of a chloroplast complex that localizes to the thylakoid membranes and contains VIPP1

Given the membrane ultrastructure defects observed under HL, we next asked where VIA1 localizes within the chloroplast and which proteins associate with it. To enable biochemical analyses, we complemented *via1* mutant cells by nuclear transformation with VIA1 genomic DNA (including endogenous regulatory regions) fused to a C-terminal 3×FLAG tag^32,33^. We confirmed VIA1–FLAG expression by immunoblotting (Fig. S5A), and complementation of the mutant phenotype was verified by HL growth assays (Fig. S5B–C). Chloroplast subfractionation showed that VIA1 is specifically enriched in the thylakoid fraction (Fig. 4A). We then performed FLAG immunoprecipitation coupled to mass spectrometry (IP–MS) under standard light and HL conditions, using untagged WT as a negative control. Under HL, VIA1–FLAG was recovered as the top hit, and VIPP1 emerged as the most prominent co-enriched interactor (Fig. 4B and Fig. S6A). Additional enriched proteins included cpUPR- and stress-associated factors, namely VIPP2 (an early and selective marker of cpUPR activation) and HSP70B (a chloroplast-localized chaperone), both previously reported as VIPP1 interactors^34,35^.

**Figure 4.**
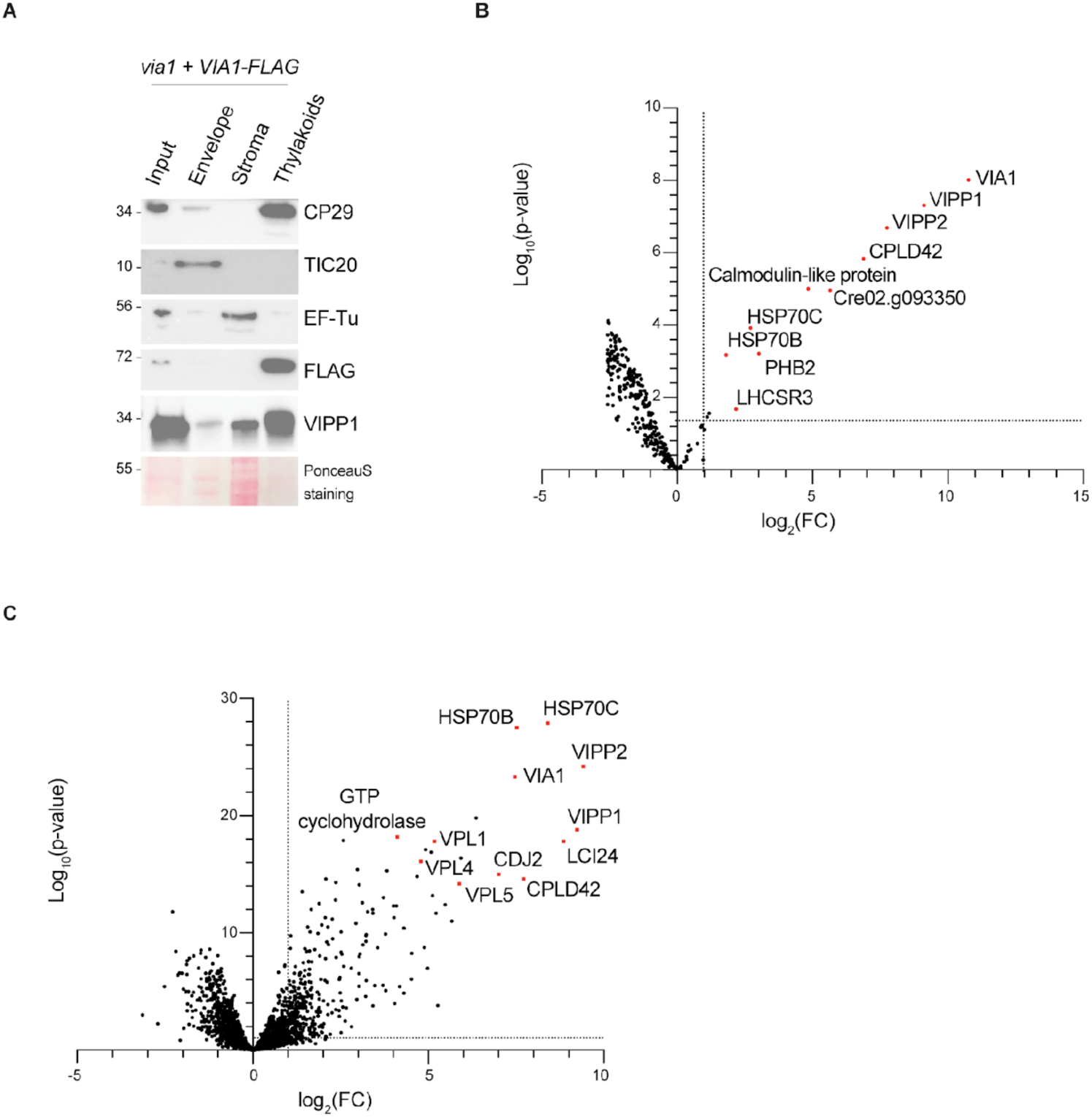
VIA1 interacts with VIPP1 and localizes to thylakoid membranes. **(A)** Chloroplast subfractionation on a two-step sucrose gradient followed by immunoblotting for VIA1–FLAG and VIPP1. CP29 (thylakoids), TIC20 (envelope), and Ef-Tu (stroma) were used as fraction markers to assess sample purity. **(B)** Volcano plot displaying log₂ fold change versus –log₁₀ p for each confidently quantified protein in FLAG co-immunoprecipitations from *via1* + VIA1-FLAG strain (three biological replicates) compared with controls (WT lysate incubated with FLAG–M2 beads). **(C)** Volcano plot of the VIPP1 co-immunoprecipitations from WT, displaying log₂ fold change versus –log₁₀ p for each confidently quantified protein. Significantly enriched proteins of interest are highlighted in red.

Considering the presence of VIPP1 in the VIA1 interactome and the similar phenotype of the *vipp1* knockdown (*vipp1-kd) a*nd the *via1* mutants, we hypothesized that VIA1 might form a complex with VIPP1 and modulate its membrane remodeling activity in stress conditions. To further test the formation of a VIA1:VIPP1 complex, we conducted the reciprocal immunoprecipitation using VIPP1 as the bait (Fig. 4C and Fig. S6B). Cell lysates from a WT Chlamydomonas strain were incubated either with protein-A Dynabeads coupled with a polyclonal anti-VIPP1 antibody or with non-coupled Dynabeads (negative control). The immunoprecipitation was conducted in normal light and HL conditions in WT and *via1* backgrounds. A total of 377 proteins were identified as potential VIPP1 interactors upon HL stress across all three technical replicates, with VIPP2 and VIPP1 being the most abundant. Heat Shock Protein 70B (HSP70B) and chloroplast DNAJ-like chaperone 2 (CDJ2) were also abundantly present. In addition, several uncharacterized proteins were detected, including VPL1–6 and VPL10, which were previously identified in a VIPP1 proximity-labeling study^35^. Importantly, peptides corresponding to VIA1 were significantly enriched in all samples, confirming the interaction between VIA1 and VIPP1. Together, these findings provide converging evidence that VIA1 and VIPP1 are binding partners *in vivo*.

### The two winged-helix domains in VIA1 mediate the interaction with VIPP1

Having established that VIA1 associates with VIPP1 *in vivo*, we next sought to define the mechanistic basis of this interaction. VIA1 is predicted to contain three transmembrane helices (DeepTMHMM 1.0^36^; Fig. 5A and Fig. S13E), consistent with its enrichment in thylakoid fractions and suggesting direct membrane anchoring. AlphaFold3 (AF3)^37^ modeling additionally identified two winged-helix domains (WH1 and WH2), a fold comprising a three-helix bundle and a four-stranded β-sheet. Although winged-helix folds are classically linked to nucleic-acid binding^38^, they are also found in ESCRT-II subunits such as Vps25^26^, where they provide protein–protein interaction surfaces for ESCRT complex assembly and ESCRT-III engagement^39,40^. This raised the possibility that VIA1 WHs might form the binding interface for VIPP1.

**Figure 5.**
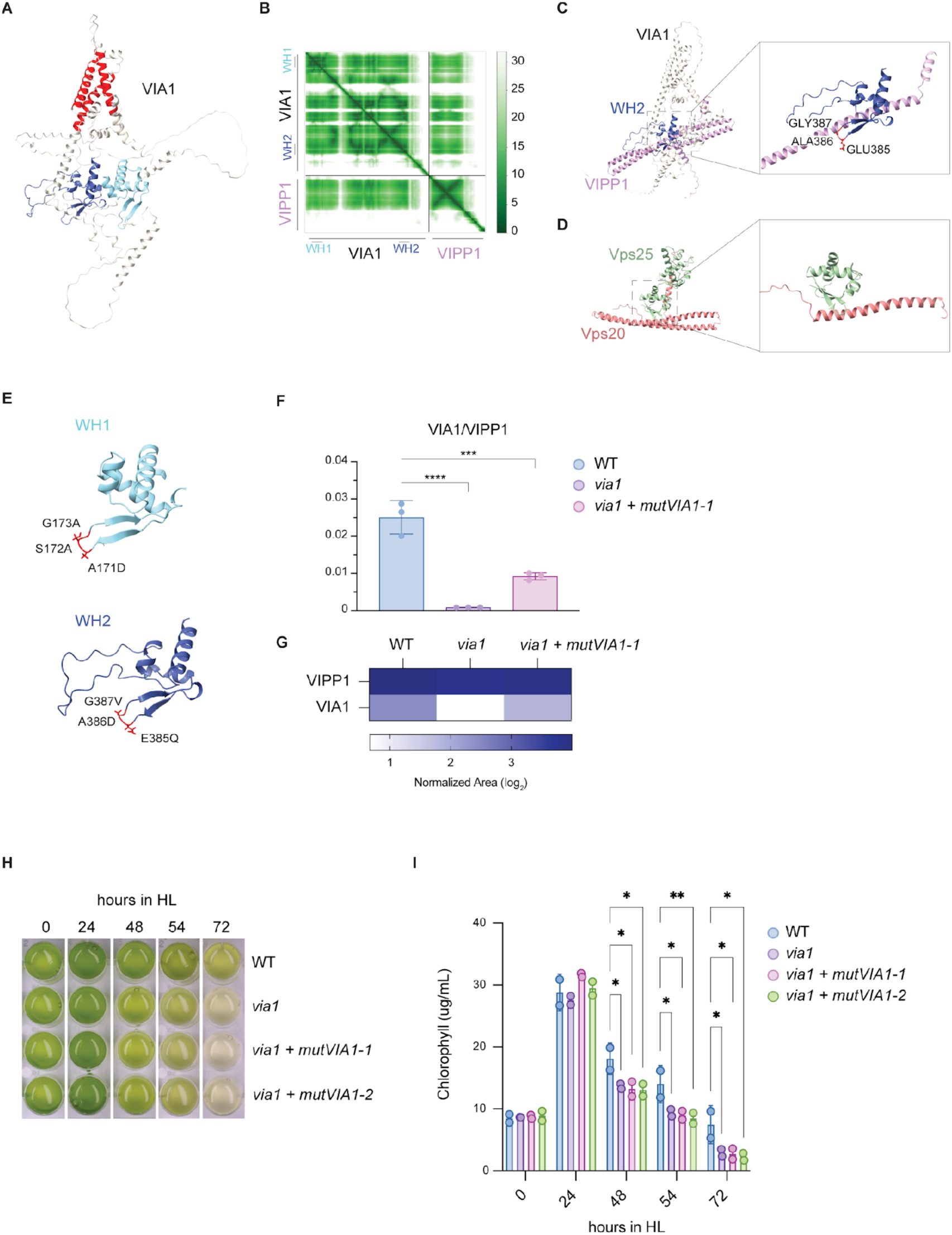
VIA1 Winged-Helix domains (WHs) mediate VIPP1 binding and VIA1 function. **(A)** AlphaFold3 (AF3) structural prediction of VIA1. WH1 and WH2 are shown in cyan and blue, respectively; the three predicted transmembrane domains are highlighted in red. **(B)** Predicted Aligned Error plot (PAE) of the AF3 prediction in (A), with WH residues indicated. **(C)** AF3 prediction of the VIA1–VIPP1 complex, with a zoom-in on the predicted interaction interface involving VIA1-WH2 and the first α-helix of VIPP1. **(D)** Crystal structure of human Vps25 and Vps20 (ESCRT-II and ESCRT-III proteins, respectively) illustrating a similar interaction interface to that predicted for VIA1 and VIPP1 (PDB 3HTU). **(E)** AF3 prediction of the two WHs showing the loop residues targeted for mutagenesis in *mutVIA1*, with the corresponding amino acid substitutions indicated. **(F)** Histogram showing the VIA1/VIPP1 abundance ratio in VIPP1 co-immunoprecipitations from WT, *via1*, and *via1* + *mutVIA1* (three technical replicates). **(G)** Heatmap of log₂ normalized areas for VIA1 and VIPP1 across the three strains used for VIPP1 co-immunoprecipitations in (F). **(H)** Representative images of liquid TAP cultures of WT, *via1*, *via1* + *mutVIA1-1*, and *via1* + *mutVIA1-2* at various time points during a 72-hour HL treatment (∼800 µmol photons m^−2^ s^−1^). *via1* + *mutVIA1-1* and *via1* + *mutVIA1-2* are two independent transformants expressing the same mutated *VIA1* transgene. **(I)** Chlorophyll measurements from the assay in (H) (three biological replicates). The significance was assessed via Two-Way ANOVA and Tukey’s multiple comparisons analysis.

Consistent with this hypothesis, AF3 prediction of the VIA1–VIPP1 complex placed the primary interface between VIA1-WH2 and the second α-helix of VIPP1 (ah2) (Fig. 5B–C). Two orthogonal analyses supported this interface: Alphabridge^41^ identified stabilizing interchain contacts enriched between VIA1-WH2 and VIPP1-ah2 (Fig. S7C), and Pickluster^42^ highlighted a recurrent interaction patch at the same region across models (Fig. S7D). Notably, in the model, the WH2 loop between the first and second β-strands was predicted to wrap around VIPP1-ah2, resembling ESCRT-II/ESCRT-III binding geometries (for example, Vps25–Vps20^39^; Fig. 5D).

To test the AF3 model, we generated a VIA1 variant (mutVIA1) by site-directed mutagenesis, introducing three point mutations in the predicted VIPP1-interacting loop of each WH domain (Fig. S7D-E). Specifically, we introduced A171D/S172A/G173A in WH1, and E385Q/A386D/G387V in WH2 (Fig. 5E) — substitutions predicted by AlphaFold3 and Alphabridge analysis to disrupt the VIA1–VIPP1 interface (Fig. S7B-C). To minimize variability in expression levels arising from random nuclear genomic integration, we inserted the mutant *VIA1* transgene into a defined neutral landing site in the chloroplast genome of *via1* cells by biolistic chloroplast transformation^43,44^, and confirmed four independent transformants by RT–qPCR (Fig. S7F).

To assess the impact of these mutations on the VIA1–VIPP1 interaction *in vivo*, we immunoprecipitated VIPP1 from WT, *via1*, and *via1* + *mutVIA1* cells. Mutant VIA1 expression was readily detected in input samples, but its co-immunoprecipitation with VIPP1 was strongly reduced compared with WT, indicating a substantially weakened association (Fig. 5F–G; Fig. S8A–B). We next asked whether disrupting the VIA1–VIPP1 interaction compromises VIA1 function. In HL growth assays, the strains expressing the binding-defective VIA1 variant failed to rescue the *via1* mutant phenotype, exhibiting reduced chlorophyll accumulation relative to WT from ∼48 h onward (Fig. 5H–I). Together, these results demonstrate that the WH domains of VIA1 are required for robust association with VIPP1 *in vivo*, and that VIA1 function in HL acclimation depends on its ability to bind VIPP1.

### Structural conservation of VIA1-WHs underlies cross-species functional complementation

As shown in Fig. 1B, VIA1 is conserved across the green lineage and is also present in cyanobacteria and diatoms. To link this sequence conservation to structural and functional conservation, we analyzed VIA1 homologs using AlphaFold structural predictions. In all species examined, VIA1 contains two WHs, one at the N terminus (WH1) and one at the C terminus (WH2) (Fig. 6A). WH1 is highly conserved across both eukaryotic and prokaryotic homologs, whereas WH2 is broadly conserved among eukaryotes but shows greater structural divergence in bacterial proteins (Fig. 6A–B). Despite this variation, the consistent presence of both WHs across lineages suggests that they are ancestral features of VIA1, likely predating plastic endosymbiosis and already present in the cyanobacterial ancestor.

**Figure 6.**
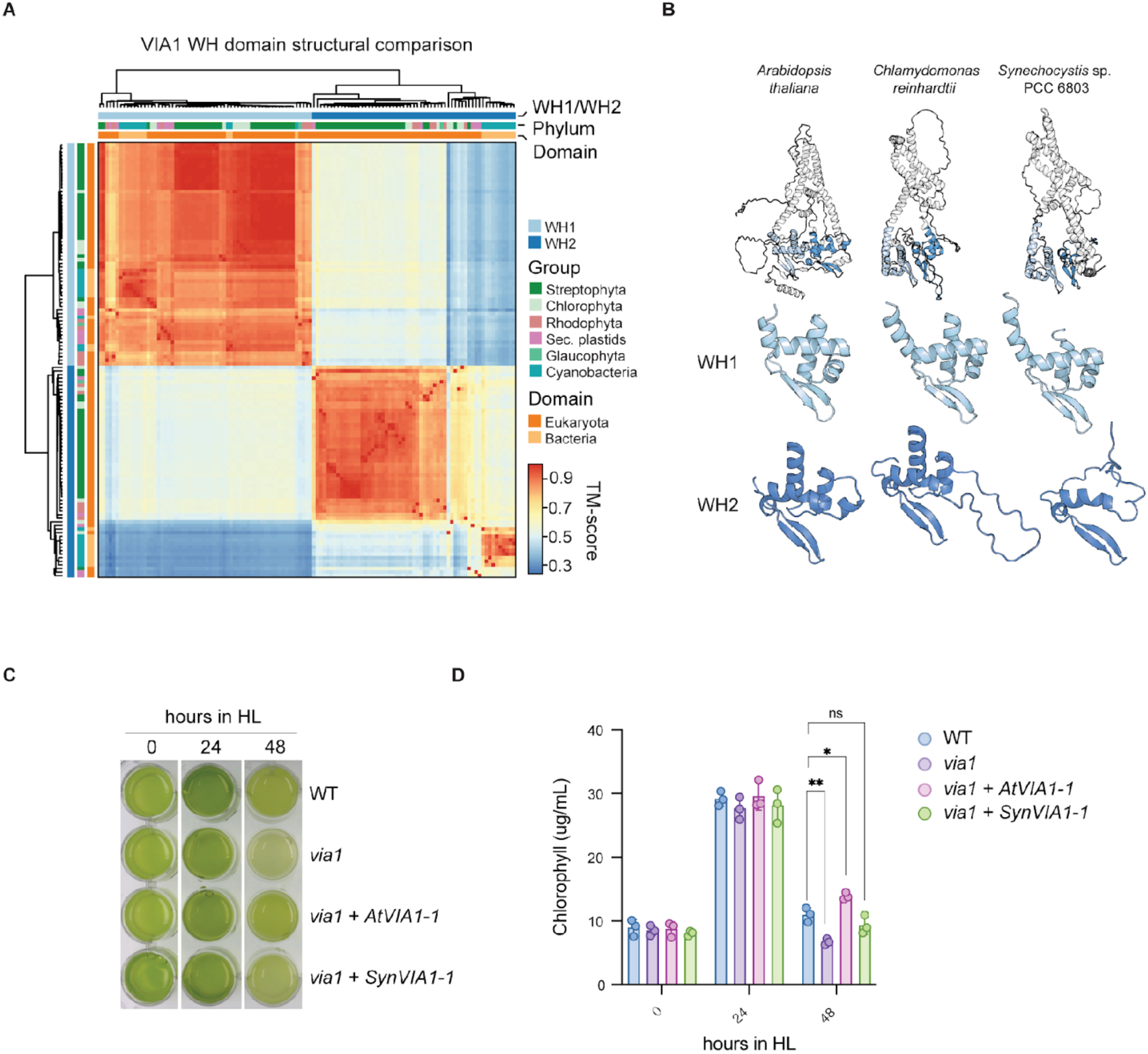
VIA1 structural conservation and cross-species complementation of the Chlamydomonas *via1.* **(A)** Structural comparison of WHs from representative eukaryotic and bacterial VIA1 proteins. Structures were predicted using AlphaFold, and WHS were defined using Chlamydomonas WH1 as a reference. Domain position and taxonomic information are noted with coloured lines. TM-score, template modeling score. **(B)** AlphaFold-predicted full-length VIA1 structures and corresponding WHs from *Arabidopsis thaliana* (UniProt ID: Q8GW20), *Chlamydomonas reinhardtii* (A0A2K3DKE5), and *Synechocystis sp.* PCC 6803 (P72990). **(C)** Representative images of liquid TAP cultures of WT, *via1*, *via1* + *AtVIA1*, and *via1* + *SynVIA1* at the indicated time points during a 48-hour HL treatment (∼800 µmol photons m^−2^ s^−1^). **(D)** Total chlorophyll content measured over the HL time course shown in (C). Statistics were performed by two-way ANOVA with Tukey’s multiple-comparisons test; ns, not significant.

Given the structural similarity between VIA1 from Chlamydomonas, Arabidopsis, and Synechocystis, we next asked whether orthologues from these distant lineages can functionally complement the Chlamydomonas *via1* mutant phenotype. To test this, we integrated untagged VIA1 orthologues from Arabidopsis (*AtVIA1*) and Synechocystis (*SynVIA1*) into the Chlamydomonas chloroplast genome of *via1* mutant cells by biolistic transformation and identified expressing strains by RT–qPCR (Fig. S9A–B).

In HL growth assays, expression of both AtVIA1 and SynVIA1 rescued the *via1* photosensitivity phenotype, with chlorophyll levels returning to near-WT values after 48 hours of HL exposure (Fig. 6C–D). Notably, SynVIA1 rescue was weaker, consistent with its greater structural divergence in WH2.

These results indicate that VIA1 performs a conserved core function, shared across cyanobacteria, algae, and land plants, that is sufficient to restore HL tolerance in Chlamydomonas.

### VIA1 interaction with VIPP1 and its role in stress protection are conserved in *Synechocystis sp.* PCC 6803

The strong structural conservation of VIA1 (Fig. 6A-B) prompted us to examine whether the VIA1–VIPP1 complex represents an evolutionarily conserved feature of photosynthetic organisms. Although this complex has been recently shown to be present in *Arabidopsis thaliana*^24^, it remained unclear whether it is also present in cyanobacteria. To test this, we generated a *Synechocystis sp.* PCC 6803 strain expressing a FLAG-tagged version of the VIA1 orthologue (SynVIA1–FLAG) (Fig. S10A), confirmed its membrane localization by fractionation (Fig. 7A–B, Fig. S10B–C) and performed FLAG immunoprecipitation followed by mass spectrometry, using an untagged strain as a negative control. SynVIA1 was the most strongly enriched protein, and VIPP1 was the top co-enriched interactor, supporting conservation of the VIA1–VIPP1 association in cyanobacteria (Fig. 7C, Fig. S11).

**Figure 7.**
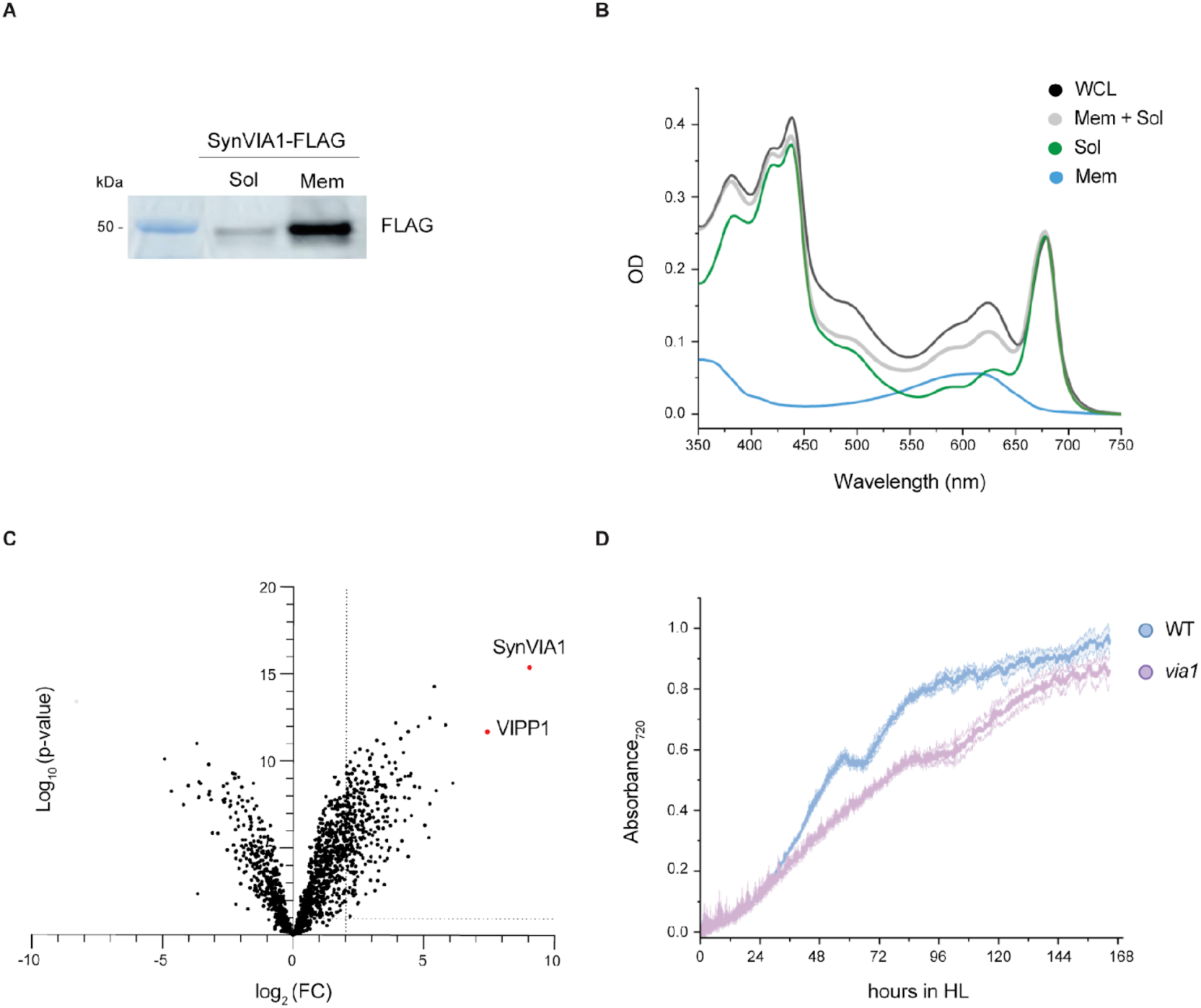
The VIA1–VIPP1 interaction and its role in stress response are conserved in *Synechocystis sp.* PCC 6803. **(A)** Whole-cell lysate (WCL) prepared from VIA1-FLAG cells was fractionated into soluble (Sol) and membrane (Mem) fractions by ultracentrifugation. A 1/10 aliquot of each fraction was subjected to FLAG immunoblot analysis. **(B)** Absorption spectra of the fractions described in (A). The WCL exhibits absorbance peaks in both the phycobilisome (PBS, associated with the soluble fraction) region (620 nm, grey dashed line) and the chlorophyll (Chl, membrane fraction) region (680 nm, grey dashed line). The soluble fraction shows a prominent peak in the PBS region with little to no signal in the Chl region, whereas the membrane fraction displays a strong Chl peak with minimal PBS signal. **(C)** Volcano plot of proteins identified by VIA1–FLAG co-immunoprecipitation mass spectrometry (IP–MS) from Synechocystis cells expressing VIA1-FLAG. Each dot represents a confidently quantified protein. The x-axis shows log_2_ fold change (FLAG IP vs. untagged WT control) and the y-axis shows −log_10_(p value). Significantly enriched proteins of interest are highlighted in red. **(D)** Growth analysis of *via1* mutant and WT cells under high light conditions (HL; 200 µmol photons m⁻² s⁻¹). Data represent the mean optical density at 720 nm (OD_720_) of four independent biological replicates, with shading indicating ± SE.

To assess whether SynVIA1 contributes to light-stress acclimation in Synechocystis, we generated an insertional *via1* mutant (Fig. S12A–B) and compared its growth with WT under normal light (NL; 80 µmol photons m⁻² s⁻¹) and high light (HL; 200 µmol photons m⁻² s⁻¹). Under NL conditions, the *via1* mutant grew similarly to WT, reaching comparable final cell densities (Fig. S12C). Under HL, however, the *via1* mutant exhibited a consistent growth defect (Fig. 7D). Whole-cell absorption spectra of HL-grown cultures further highlighted differences between the two strains: whereas WT and *via1* cells showed similar absorption in the chlorophyll and phycobilisomes regions (∼675 nm and ∼630 nm, respectively), *via1* cells displayed reduced absorption in the carotenoid region (460–520 nm), suggesting an impaired HL acclimation response (Fig S12D).

## Discussion

Thylakoid membrane integrity is fundamental to photosynthetic function yet can be compromised, especially under saturating light conditions, when protein and lipid damage must be continuously repaired. VIPP1, an ESCRT-III-like protein, is one of the most intensively studied and highly conserved factors implicated in thylakoid biogenesis and maintenance across both bacterial and eukaryotic photosynthetic lineages. Here, by leveraging the chloroplast unfolded protein response (cpUPR)—a signaling pathway triggered by proteotoxic and membrane stress—we identified VIA1 as a previously uncharacterized factor that interacts with VIPP1. In contrast to VIPP1, however, VIA1 function becomes most evident during high light stress, consistent with a role in stress-responsive membrane remodeling rather than in constitutive membrane biogenesis.

Structurally, VIA1 contains two conserved winged-helix domains and engages the α1 helix of VIPP1 through an interface that is functionally required. This interaction architecture is conceptually reminiscent of ESCRT-II/ESCRT-III contacts that coordinate membrane remodeling in eukaryotic cells^45^. However, VIA1 differs fundamentally from canonical ESCRT-II in both organization and apparent function. Eukaryotic ESCRT-II is a soluble, hetero-tetrameric cytosolic complex that assembles as a Y-shaped scaffold and promotes ESCRT-III recruitment and polymer formation at membranes^40,46^. In contrast, VIA1 is a membrane-embedded protein. This imposes a distinct topological constraint: if both WH domains engage VIPP1 from the same side of the membrane, VIA1 would require an additional membrane-spanning segment not evident in the current AlphaFold model. As shown in Fig. S13, a conserved hydrophobic stretch between the first transmembrane helix and the second WH domain is a plausible candidate for this role in VIA1, though the presence of charged residues makes it atypical of canonical transmembrane helices. Resolving the membrane topology of VIA1 and its precise mode of engagement with VIPP1 will be an important goal for future structural and biochemical studies.

The WH domains mediating VIPP1 interaction are ancestral features of VIA1, present in both prokaryotic and eukaryotic homologs, suggesting that the VIA1–VIPP1 interaction module predates plastid endosymbiosis. Consistent with this, VIA1 orthologs from Arabidopsis and Synechocystis partially rescue the Chlamydomonas *via1* mutant phenotype, whereas disruption of VIA1 in Synechocystis compromises growth under light stress, mirroring the functional requirement observed in Chlamydomonas. These results are further supported by recent work showing that Chlamydomonas VIA1 can functionally complement the Arabidopsis *via1* mutant^24^.

Together, these findings support the existence of an evolutionarily conserved VIA1–VIPP1 complex involved in thylakoid membrane remodeling that likely predates plastid endosymbiosis, advancing our understanding of membrane homeostasis in photosynthetic organisms.

## Author Contributions

S.R. conceived the project, supervised Chlamydomonas research activities, provided resources, and coordinated collaborations. P.V. performed the majority of experiments, analyzed data, carried out structural predictions, curated data, and prepared figures, with input from all co-authors. L.M. assisted P.V. with the cloning of selected constructs and contributed to high light growth assays. E.T. performed most photosynthetic measurements and analyzed and curated the related data, under the supervision of T.M. K.K. generated the Synechocystis mutant and tagged strains and conducted all Synechocystis-related experiments, under the supervision of Y.M., except for the co-Immunoprecipitation–mass spectrometry analysis, performed by P.V. N.A.T.I. carried out phylogenetic analyses and visualized the corresponding results. S.R. and P.V. wrote and edited the manuscript with input from all co-authors. All authors reviewed and approved the final manuscript.

## Acknowledgments

We thank the Protein Chemistry Facility at IMP/IMBA/GMI for mass spectrometry measurements and proteomics data analysis performed using the Vienna BioCenter Core Facilities (VBCF) instrument pool. We are grateful to Nicole Drexter (VBCF) for assistance with electron microscopy, and to Sven Klumpe and members of the Ramundo lab for valuable discussions. We also thank Natalie Page and J. Matthew Watson for editing and proofreading, and Liam Dolan, Youssef Belkhadir, and the anonymous PNAS reviewers of the previous manuscript version for critical comments.

## Materials and Methods

### Growth conditions (*C. reinhardtii*)

*Chlamydomonas reinhardtii* strains were maintained on solid Tris-Acetate-Phosphate (TAP) media containing 1.6% [w/v] agar (USP grade, Thermo Fisher Scientific) and Hutner’s trace elements (Chlamydomonas Resource Center) at 22 °C under low light conditions (∼30 µmol photons m^−2^ s^−1^). CLiP mutant strains harboring the CIB1 insertion cassette encoding the *APHVIII* paromomycin resistance marker (CrRS432, CrRS433; see Table S2 for details) were maintained on solid TAP media supplemented with 10 µg mL^−1^ paromomycin. Unless otherwise stated, strains were grown in liquid culture under standard light conditions with an intensity of 80 µmol photons m⁻² s⁻¹ and in the absence of any antibiotics. A complete list of strains used in this research can be found in Table S2.

### Growth conditions (*Synechocystis sp.* PCC 6803)

Cyanobacteria were cultured in BG11 medium supplemented with 6 µg/ml Ferric ammonium citrate, and 5 mM glucose, continuous white light (8 µmol photons m^−2^ sec^−1^) provided by T5 fluorescent lights at 26 °C, shaken at 120 rpm. Before harvesting cells for MS analysis, they were adapted to a light intensity of 20 µmol photons m^−2^ sec^−1^ for five days.

### Growth assay (*Synechocystis sp.* pcc 6803)

Cells were initially grown for 14 days in 50 mL flasks of BG11 supplemented with 6 µg/ml Ferric ammonium citrate. Cultures were then diluted into 500 ml flasks and acclimated to low light intensity (8 µmol photons m^−2^ sec^−1^) for three days before inoculation into the bioreactor (Multi-Cultivator MC 1000-OD, Photon Systems Instruments). After this acclimation period, each culture was split into four 50mL subcultures, which were used to initiate growth measurements in the bioreactor. Light intensity was set to either normal light (80 µmol photons m^−2^ sec^−1^) or high light (200 µmol photons m^−2^ sec^−1^), and absorbance at both 720 nm and 680 nm was recorded every 10 minutes. Cultures were bubbled with air throughout growth. Fluorescence emission spectra were recorded on a Fluoromax-4 spectrofluorometer (HORIBA Jobin-Yvon). For 77 K measurements, slit width of 5 nm and 3 nm were used for the entrance and exit monochromators, respectively. Samples were diluted to an OD_680_ 0.1 in a final glycerol concentration of 40%, frozen around thin glass rods in liquid nitrogen, and excited at 440 nm. Whole cell absorbance was measured after 180 h of bioreactor growth using a Cary 4,000 UV–Vis spectrophotometer (Agilent Technologies) equipped with an integration sphere.

### Isolation and genotyping of *C. reinhardtii* VIA1 insertion strains

Genomic DNA extraction from WT, *via1-1,* and *via1-2* liquid cultures in the exponential growth phase and in standard light conditions (∼80 µmol photons m^−2^ s−^1^) was carried out as previously described^23^. The insertion of the mutagenic cassette in the VIA1 locus was verified by PCR in each mutant. All PCRs were performed using KOD Hot Start DNA Polymerase (EMD Millipore). In the *via-1* strain, the insertion occurs in exon 10. Therefore, primer pairs oRS39 and oRS40 were designed to anneal in exon 9 and the 3’UTR, respectively (see Fig. S2 for details). This primer pair allows the amplification of the WT locus as well as of the mutated locus containing the cassette (PCR conditions: 95 °C 2 min, 95 °C 20 sec, 55 °C 10 sec, 70 °C 1 min 40 sec). To selectively amplify a section of the cassette inserted in the *CrVIA1* locus, the oRS01 primer was designed to anneal within the mutagenic cassette and was used in conjunction with oRS39 (PCR conditions: 95 °C 2 min, 95 °C 20 sec, 55 °C 10 sec, 70 °C 28 sec). For *via-2*, which harbors the insertion within intron 7, primer pairs oRS37 and oRS38 were designed to anneal in exon 6 and exon 8, respectively. This primer pair allows the amplification of the WT locus as well as of the mutated locus containing the mutagenic cassette (PCR conditions: 95 °C 2 min, 95 °C 20 sec, 55 °C 10 sec, 70 °C 1 min 40 sec). Moreover, to selectively amplify a section of the cassette inserted in the *CrVIA1* locus, the oRS30 primer was designed to anneal within the mutagenic cassette and was used in conjunction with oRS38 (PCR conditions: 95 °C 2 min, 95 °C 20 sec, 55 °C 10 sec, 70 °C 30 sec). To ensure the quality of the employed genomic DNA (gDNA), a mating type-specific PCR^47^ was performed as a control. All PCR reactions were subjected to electrophoresis on 1% [w/v] agarose gels to verify their expected sizes.

### Backcross and tetrad dissection analysis

Cells were streaked onto a fresh TAP agar plate and grown under light (80 µmol photons m^−2^ s^−1^) for approximately 3-4 days. The cells were subsequently restreaked onto a TAP agar plate containing one-tenth of the standard NH_4_Cl concentration to induce nitrogen starvation and gametogenesis. The day after, gametes were suspended in 100 µL of sterile water to a final density of ∼1 × 10⁶ cells mL⁻¹ and incubated with gentle agitation for 30-60 minutes under light (80 µmol photons m^−2^ s^−1^). Mating efficiency was monitored every 30 minutes by mixing 2 µL of the mating mixture with 8 µL of Lugol’s solution (Gatt-Koller) and scoring quadriflagellate formation under a microscope. Mating typically occurred within 2 h. Subsequently, 100 µL of the mating mixture was spotted onto the center of a TAP 4% [w/v] agar plate and incubated in the dark (covered with aluminum foil) for 6 days to allow zygote maturation. Vegetative cells were then gently removed by scraping the plate surface with a rectangular razor blade (Apollo Herkenrath), leaving zygotes attached to the agar. To recover zygotes, 100 µL of liquid TAP was added to the plate surface, and zygotes were detached with a different razor blade (Aesculap AG). The suspension was transferred onto several fresh TAP plates containing 1.6% [w/v] agar and spread in a line in the middle of each plate. Plates were then incubated under light (80 µmol photons m^−2^ s^−1^) for 24 h to induce germination. Tetrads were identified and dissected using a SporePlay+ dissection microscope (Singer Instruments). Once colonies became visible to the naked eye, progeny were re-arrayed into a 96-format grid and replicated onto TAP plates with or without paromomycin for segregation analysis.

### RNA extraction and RT-qPCR for gene expression analysis (*C. reinhardtii*)

Total RNA was extracted from WT, *via1-1*, and *via1-2* liquid cultures in exponential phase grown under standard light (80 µmol photons m^−2^ s^−1^) as described previously^23^. For cDNA synthesis, 3 µg total RNA and oligo(dT)₁₈ were used with the Maxima First Strand cDNA Synthesis Kit (Thermo Fisher Scientific) following the manufacturer’s instructions. cDNA was diluted 1:10 prior to quantitative PCR. qPCR was performed using SYBR Green chemistry (Bio-Rad) according to the manufacturer’s instructions. Primer sequences used to amplify VIA1 are provided in Table S5. No-template controls (master mix without cDNA) were included to assess contamination and non-specific amplification. Ct values were analyzed following the “eleven golden rules”, as described previously^48^. Transcript abundance was normalized to the reference gene *GBLP* (Cre06.g278222). Two biological replicates were analyzed per strain, each measured in two technical replicates.

### Phylogenetics and species distributions

To reconstruct the VIA1 phylogeny, the *C. reinhardtii* ortholog (Cre07.g338350) was searched against a series of predicted proteomes representing eukaryotic and prokaryotic diversity using Diamond BLASTP v2.1.11 (ultra-sensitive, E < 10^−5^)^49^. These datasets comprised predicted chlorophyte proteomes (*n =* 13), plant proteomes (*n* = 37), the EukProt v3 TCS dataset, and selected prokaryotic reference proteomes (*n =* 5,143) from UniProt^50,51^. The resulting hits were extracted and aligned with MAFFT v7.520, and alignments were trimmed with a gap-threshold of 50% using trimAl v1.4^52,53^. Initial phylogenies were inferred using IQ-Tree v2.3.6 and the LG4M substitution model with support derived from Shimodaira-Hasegawa approximate likelihood ratio tests (SH-aLRT, *n* = 1,000)^54,55^. Orthologs were selected phylogenetically and extracted before being re-aligned with MAFFT using the L-INS-i algorithm. The alignment was trimmed with a gap-threshold of 10% and used to generate a hidden Markov model (HMM) using HMMER v3.1b2^56^. To increase search sensitivity, proteomes were re-searched using the VIA1 HMM (E < 10^−5^) and the resulting hits were curated phylogenetically as described. To generate the final phylogeny, an alignment was generated using VIA1 homologs and MAFFT L-INS-i before being trimmed with a gap-threshold of 50%. The phylogeny was inferred using IQ-Tree and the LG+C60+F+I+R8 substitution model selected using ModelFinder^57^. The phylogeny was visualized using FigTree v1.4 (http://tree.bio.ed.ac.uk/software/figtree/) and ITOL v6^58^. The curated VIA1 homologs were further used to assess the distribution of VIA1 across photosynthetic species. To account for missing gene models, the genomes of taxa lacking VIA1 were searched using MiniProt v0.18, and homologs were predicted and phylogenetically curated when found^59^. Protein sequences, alignments, phylogenies, and datasets are available from here: https://figshare.com/s/c411a207630c1d30d379

### Structural comparisons

To assess structural conservation in the winged helix domains across VIA1 homologs, predicted protein structures were generated using AlphaFold2 and the MMseqs2 database (*n* = 62)^37,60^. The highest-ranking model (*n* = 5) for each protein was selected and split at its mid-point, generating N- and C-terminal halves. To extract the winged helix (WH) domains, the N-terminal WH from *C. reinhardtii* was isolated in PyMOL v2.5.0 (residues 35-98) and aligned to both the N- and C-terminal regions of each predicted VIA1 structure using TM-align v20220412 (query coverage > 50%)^61^. Aligned regions were extracted, and pairwise TM-scores were calculated using TM-align. Predicted protein structures were visualized in PyMOL.

### Total protein extraction and immunoblot analysis (*C. reinhardtii*)

For all experiments, total protein extracts were prepared from whole-cell lysates under denaturing conditions. Cells from 5 mL cultures in the exponential growth phase were harvested by centrifugation at 3,000 × g for 5 min, and pellets were resuspended in 150 µL SDS lysis buffer (100 mM Tris-HCl, pH 8.0, 600 mM NaCl, 4% [w/v] SDS, 20 mM EDTA) freshly supplemented with EDTA-free protease inhibitor cocktail (Roche). Samples were vortexed for 10 min at room temperature, incubated for 30 min at 37 °C, and clarified by centrifugation at 21,300 × g for 15 min at room temperature to remove insoluble debris.

The supernatant was transferred to a new tube and supplemented with ¼ volume of 5× SDS loading buffer (250 mM Tris-HCl, pH 6.8, 5% [w/v] SDS, 0.025% [w/v] bromophenol blue, 25% [v/v] glycerol), freshly supplemented with 5% [v/v] 2-mercaptoethanol. A 5 µL aliquot of each extract was retained for protein quantification using a bicinchoninic acid (BCA) assay according to the manufacturer’s instructions (Sigma-Aldrich). Specifically, 2 µL of protein extract was mixed with 200 µL BCA working solution (1:50 BCA: copper sulfate), incubated for 5 min at 50 °C, and absorbance was measured at 562 nm using a BSA standard curve.

For immunoblotting, 40 µg of total protein per sample was resolved by SDS–PAGE using Criterion precast gels (Bio-Rad) and transferred onto 0.2 µm nitrocellulose membranes (Amersham). Membranes were blocked for 1 h at room temperature in PBS-T supplemented with 5% [w/v] nonfat dry milk (Maresi). Primary and secondary antibodies (listed in Table S1) were diluted in blocking buffer. Primary antibody incubations were performed for 1 h at room temperature, except for anti-D1 and anti-VIPP2, which were incubated overnight at 4 °C. HRP-conjugated anti-rabbit or anti-mouse secondary antibodies (Promega) were applied at 1:10,000 for 1 h at room temperature.

Following each antibody incubation, membranes were washed three times for 10 min in PBS-T containing 1% [w/v] nonfat dry milk. Signals were detected using luminol-based enhanced chemiluminescence (SuperSignal West Dura or West Femto; Thermo Fisher Scientific).

### Immunoblot analysis (*Synechocystis sp.* PCC 6803)

For immunoblotting, 100 µg of total protein per sample was resolved by SDS–PAGE using Criterion precast gels (Bio-Rad) and transferred onto 0.45 µm nitrocellulose membranes (GVS #S74181100). Membranes were blocked overnight at 4 °C in PBS-T supplemented with 5% [w/v] nonfat dry milk (Nestle Carnation). Primary antibodies (listed in Table S1) were diluted in blocking buffer. For Flag detection an HRP conjugated OctA-Probe Antibody from Santa Cruz Biotechnology. Signals were detected using luminol-based enhanced chemiluminescence (West Femto; Thermo Fisher Scientific).

### Chlorophyll Fluorescence Measurements (*C. reinhardtii*)

Chlorophyll fluorescence measurements of *Chlamydomonas* were performed using a Dual PAM 100 fluorometer (Walz, Germany). Cells were grown in minimum media for 5 days, harvested during the exponential growth phase and subsequently subjected to a 30-minute period of dark adaptation while maintaining gentle agitation. The following parameters were calculated: Photosystem I Quantum Yield ((Pm - Pm’)/Pm), Photosystem II Quantum Yield (Fv/Fm) and Non-Photochemical Quenching ((Fm - Fm’)/Fm’) where Fm’ is the maximal chlorophyll fluorescence yield during exposure to light. Calculations were based on established methodologies for the measurement of photosynthetic parameters^62,63^. New measurements in stress conditions were performed using an AquaPen AP110/C fluorometer (Photon Systems Instruments). Maximum quantum yield of PSII (Fv/Fm) was measured in liquid cultures over a 48-hour growth assay under normal light (80 µmol photons m⁻² s⁻¹) and high light (800 µmol photons m⁻² s⁻¹). Experimental conditions, including cell density and chlorophyll concentration at the start of the assay, are described in the “High light assay and time course” section. Fv/Fm (Qy) measurements were performed using the blue measuring light (455 nm) with a 30% flash pulse, a 70% super pulse, and an actinic pulse of 300 µmol photons m⁻² s⁻¹.

### Spot growth assay

Liquid cultures were initiated from freshly streaked colonies (3–4 days old) grown on TAP agar under continuous light (80 µmol photons m⁻² s⁻¹). Cells were inoculated into 15 mL TAP in 100 mL glass flasks and grown for 3 days under continuous light (80 µmol photons m⁻² s⁻¹) with shaking at 130 rpm. Cultures were then expanded to 50 mL and grown for an additional day. Cell densities were determined and cultures were normalized to 1 × 10⁶ cells mL⁻¹, followed by 10-fold serial dilutions to 1 × 10⁵, 1 × 10⁴, and 1 × 10³ cells mL⁻¹. For each strain, 4 µL of each dilution was spotted onto TAP agar plates. Plates were incubated under standard light (80 µmol photons m⁻² s⁻¹), and images were acquired after 4 and 7 days.

### High light assay and time course (*C. reinhardtii*)

Liquid cultures were initiated from freshly streaked colonies (3-4 days old) grown on TAP plates under continuous light (80 µmol photons m^−2^ s^−1^). Cells were inoculated into 15 mL TAP in 100 mL glass flasks and grown for 2–3 days under continuous light (80 µmol photons m^−2^ s^−1^) with shaking at 130 rpm. Once cultures reached ∼1 × 10⁶ cells mL⁻¹, they were expanded to 50 mL and grown for an additional 1–2 days. Total chlorophyll was quantified by methanol extraction as described previously¹⁷, and cultures were normalized to 8 µg chlorophyll mL⁻¹ in a final volume of 31 mL in 50 mL Falcon tubes. Three technical replicates were prepared for each strain. The high light treatment was performed at ∼800 µmol photons m^−2^ s^−1^, employing a Plant LED Grow Light (Phlizon, Cob Series 3000W) positioned ∼40 cm away from the cultures. Light intensity was measured along the length of each tube to ensure uniform illumination, and tubes were randomly repositioned every 24 h to minimize positional effects. Transfer to HL conditions resulted in a temperature increase of ∼4–5 °C.

For HL growth assays, cultures were photographed, and chlorophyll content was measured at the time points indicated for each experiment. For the HL time-course immunoblot experiment, cultures were adjusted to a final volume of 100 mL at the start of HL exposure and illuminated as above for 8 h. 5 mL aliquots were collected at 2-h intervals, pelleted by centrifugation (3,000 × g, 5 min), and stored at –20 °C for subsequent total protein extraction.

### Transmission electron microscopy (*C. reinhardtii*)

Wild-type (WT) and *via1-2* cell cultures were subjected to HL (1000 µmol photons m^−1^ s^−1^) for 0, 2, 4, and 6 h. 10 mL aliquots from each time point, at a concentration of approximately 1 × 10^6^ cells mL^−1^, were gently spun down at 500 x g for 3 minutes, and the resulting cell pellets were fixed using a mixture of 2% [v/v] paraformaldehyde (Electron Microscopy Sciences, Hatfield, PA) and 2% [v/v] glutaraldehyde (Agar Scientific, Essex, UK) in 0.1 M sodium cacodylate buffer, pH 7.4. This fixation process was carried out overnight at room temperature. Samples were then rinsed with the same buffer, post-fixed in 2% [w/v] osmium tetroxide (Agar Scientific, Essex, UK) in 0.1 M sodium cacodylate buffer, pH 7.4, dehydrated in a graded series of acetone, and embedded in Agar 100 resin (Agar Scientific, Essex, UK). Subsequently, 70 nm sections were cut and post-stained with 2% uranyl acetate [w/v] and Reynolds lead citrate (Delta Microscopies, Mauressac, France). Micrographs were recorded on a FEI Morgagni 268D (FEI, Eindhoven, The Netherlands) operated at 80 keV, equipped with a Mega View III CCD camera (Olympus-SIS).

### Chloroplast subfractionation

Chloroplast subfractionation was performed using a protocol modified from^64^. *via1-1* + VIA1 cultures were grown in liquid TAP under continuous light (80 µmol photons m⁻² s⁻¹) for 3 days to a density of ∼1 × 10⁶ cells mL⁻¹. Cells were harvested by centrifugation and resuspended in an equal volume of 0.33 MS buffer (25 mM MES-NaOH pH 6.5, 1.5 mM NaCl, 0.33 M sucrose) supplemented immediately before use with EDTA-free protease inhibitor cocktail (Roche). Cells were lysed by five freeze–thaw cycles (snap-freezing in liquid nitrogen and thawing at room temperature).

Subcellular fractionation was performed in open-top polyclear ultracentrifuge tubes (SETON Scientific, 7010) containing a two-step sucrose gradient prepared in 0.33 MS buffer (2 mL 1.3 M sucrose over 2 mL 1.8 M sucrose). 200 µL lysate was layered onto the 1.3 M cushion and centrifuged at 139,863 × g for 30 min at 4 °C (SW60 Ti rotor; Beckman Coulter). The supernatant above the 1.3 M cushion (stroma fraction) was collected, transferred to polycarbonate tubes (Beckman Coulter, 343775), and clarified at 150,000 × g for 1 h at 4 °C (TLA-100 rotor; Beckman Coulter) to remove residual debris.

Envelope membranes migrated as a yellow/orange band at the interface between the stromal fraction and the 1.3 M sucrose cushion. The envelope fraction (∼150 µL) was collected and pelleted at 150,000 × g for 1 h at 4 °C. The pellet was resuspended in 150 µL 0.33 MS buffer and re-pelleted at 150,000 × g for 30 min at 4 °C. The final envelope pellet was resuspended in 50 µL 0.33 MS buffer.

Thylakoid membranes accumulated at the interface between the 1.3 M and 1.8 M sucrose cushions. The thylakoid band (∼200 µL) was collected, diluted 1:1 with 0.33 MS buffer, and pelleted at 15,000 × g for 15 min at 4 °C in a tabletop centrifuge.

Protein concentrations were determined by BCA assay using 5 µL aliquots from each fraction. Fractions were mixed with 5× SDS loading buffer (final 1×; ¼ volume added), incubated for 5 min at 95 °C, and analyzed by SDS–PAGE followed by immunoblotting. Compartment-specific marker proteins were used to assess fraction purity (see Results). Details of loading buffer composition, BCA assay conditions, and immunoblotting procedures are provided in the corresponding sections above.

### Membrane subfractionation (*Synechocystis sp.* PCC 6803)

FLAG-VIA1 cells were grown in 500 mL of BG11 medium supplemented with Fe and glucose for 6 weeks. Cells were harvested by centrifugation at 12,000 rpm (18,514 x g) for 3 min and resuspended in STN1 buffer (30 mM Tricine pH8, 0.4 M sucrose, 15 mM NaCl). DNase (5 µL) was added, and cells were lysed by passage through a French press (Constant systems CF1) at 30,000 psi for three cycles. The lysate was centrifuged at 12,000 rpm (18,514 x g) for 5 min to remove cellular debris, the supernatant was termed the whole cell lysate fraction (WCL). The WCL was subjected to ultracentrifugation at 45,000 rpm (208,000 x g) for 3 h at 4 °C. The supernatant was collected as the S1 fraction (the volume of this fraction was the reference volume for all washes and resuspension step). The S1 fraction was subjected to three additional ultracentrifugation steps at 45,000 rpm (208,000 x g) for 3 h at 4 °C to yield the final soluble fraction (Sol). The membrane pellet of the first ultracentrifugation step was resuspended in 0.75 volume of high-salt STN1 buffer (150 mM NaCl) and ultracentrifuged again at 45,000 rpm (208,000 x g) for 3 h at 4 °C. The pellet was resuspended again in 0.75 volume STN1 buffer to generate the membrane (Mem) fraction.

### Co-Immunoprecipitation (*C. reinhardtii*)

A 5 mL preculture of *C. reinhardtii* WT cells (CrRS166) was grown in Tris–Acetate–Phosphate (TAP) medium at 22 °C under standard light conditions until a density of 2–6 × 10⁶ cells mL⁻¹ was reached. Cultures were sequentially expanded to 50 mL and then 150 mL to reach a final density of 2-6 × 10⁶ cells mL⁻¹, at which point cells were harvested by centrifugation (3,000 × g, 5 min, 4 °C). The pellet was resuspended in a 1:1 (v/v) ratio of cell pellet to ice-cold 2× IP buffer (100 mM HEPES, 100 mM potassium acetate, 4 mM magnesium acetate, 2 mM calcium chloride, 400 mM sorbitol) supplemented immediately before use with EDTA-free protease inhibitors (Roche). Samples were snap-frozen in liquid nitrogen.

Cells were lysed by five freeze–thaw cycles (alternating thawing at room temperature and snap-freezing in liquid nitrogen). For solubilization, 300 µL lysate was mixed with 300 µL ice-cold IP buffer and 66 µL of 10% (w/v) digitonin (EMD Millipore) to achieve a final digitonin concentration of 1% (w/v). Samples were incubated on a rotating wheel (1 h, 4 °C) and clarified by centrifugation (21,300 × g, 30 min, 4 °C).

For immunoprecipitation, 25 µL Protein A Dynabeads (Thermo Fisher) were used per sample. Beads were washed four times with 1 mL IP buffer, resuspended in the original volume, and incubated with 4 µL of 1 mg mL⁻¹ anti-VIPP1 antibody per sample (45 min, 20 °C, rotation). As negative controls, lysates were incubated with non-antibody-coupled beads.

After magnetic separation and one wash with IP buffer, beads were washed twice with IP buffer containing 0.1% (w/v) digitonin (5 min, 4 °C). Subsequently, 250 µL clarified lysate was added to the antibody-bound beads and incubated (3 h, 4 °C, rotation). Beads were then washed four times with IP buffer + 0.1% digitonin (first two washes 5 min; final two washes 10 min; all at 4 °C).

Elution was performed under denaturing conditions by resuspending beads in 60 µL 2× denaturing lysis buffer (100 mM Tris-HCl pH 8.0, 600 mM NaCl, 4% SDS, 20 mM EDTA) and incubating at 70 °C for 15 min (850 rpm). After magnetic separation, the supernatant was collected and stored at –20 °C.

For mass spectrometry preparation, 20 µL eluate was mixed with 5 µL 5× SDS loading buffer (250 mM Tris-HCl pH 6.8, 5% SDS, 0.025% bromophenol blue, 25% glycerol) supplemented with 5% (v/v) β-mercaptoethanol, incubated 5 min at 95 °C, separated by SDS-PAGE, and stained with colloidal Coomassie (MBS, VBCF). Gel lanes were excised, placed in 1% acetic acid, and stored at 4 °C until processing.

The same procedure was used for FLAG co-immunoprecipitation in *C. reinhardtii*, except that Magnetic Anti-FLAG M2 beads (Sigma-Aldrich) were used (25 µL per sample), and no antibody coupling step was required. The elution step was carried out under native conditions: beads were resuspended in 35 uL of 2X IP buffer supplemented with 10% digitonin and 1 μg/uL of 3XFLAG peptide (Peptide Facility, VBC), and incubated at 4 °C for 30 min in a thermomixer at 850 rpm. The supernatant was then separated and stored at −20 °C. This elution step was repeated 3 times, and the three elutions were pooled.

### Co-Immunoprecipitation (*Synechocystis sp.* pcc 6803)

*Synechocystis* cells expressing VIA1 with an endogenous C-terminal FLAG tag were grown on selective BG11 agar plates for two weeks under continuous white light (∼8 µmol photons m⁻² s⁻¹). Colonies were inoculated into 50 mL BG11 and grown for two weeks, then expanded to 500 mL BG11 for two additional weeks. Cultures were further scaled to 10 L BG11 and grown for one week. For light adaptation, 500 mL cultures were exposed to 20 µmol photons m⁻² s⁻¹ for two weeks prior to harvest. Cells were collected by centrifugation and resuspended in an equal volume of ice-cold 2× IP buffer (see previous paragraph). The suspension was added dropwise into liquid nitrogen to generate frozen pellets, which were homogenized using a pre-chilled cryo-mill (Retsch; 5 cycles of 3 min at 20 s⁻¹, with 2 min cooling intervals in liquid nitrogen). The powdered material was thawed on ice and further homogenized by 30 strokes in a Kontes Dounce homogenizer.

For solubilization, 300 µL lysate was mixed with 300 µL IP buffer and 66 µL 10% digitonin (final concentration 1%), incubated 1 h at 4 °C, and clarified by centrifugation (21,300 × g, 30 min, 4 °C). From this point on, FLAG immunoprecipitation was performed as described in the previous paragraph.

### Processing of Mass-Spectrometry samples (in-gel digest)

Coomassie-stained gel bands were cut to 2-3 mm pieces, transferred to 0.6 ml tubes and incubated with different solutions by shaking for 10 minutes at room temperature followed by removal of the supernatant as follows: Gel pieces were washed with 200 µl 100 mM ammonium bicarbonate (ABC), destained by two repeated rounds of shrinking in 200 µl 50% ACN in 50mM ABC and reswelling in 200 µl 100 mM ABC. Gel pieces were shrunk with 100 ul ACN before being reduced with 100 µl of 10 mM Dithiothreitol in 100 mM ABC by incubation at 57 °C for 30 min and alkylated with 100 µl of 20 mM Iodoacetamide in 100 mM ABC by incubation at RT for 30 min.

Wash steps were repeated as described for destaining, and gel pieces were shortly dried in a speed-vac after the final shrinking step. Gel pieces were soaked in 12.5ng/ul Trypsin in ABC for 5min at 4 °C. Excess solution was removed, ABC was added to cover the pieces, and samples were kept overnight at 37 °C. The supernatant containing tryptic peptides was transferred to a fresh tube, and gel pieces were extracted by the addition of 20 µL 5% formic acid and sonication for 10 min in a cooled ultrasonic bath. This step was performed twice. All supernatants were unified. A similar aliquot of each digest was analysed by LC-MS/MS.

### NanoLC-MS/MS Analysis

The nano HPLC system (Vanquish Neo UHPLC-System, Thermo Scientific) was coupled to an Orbitrap Exploris 480 mass spectrometer equipped with a FAIMS pro interface or to an Orbitrap Astral mass spectrometer, both equipped with a Nanospray Flex ion source (Thermo Scientific). Peptides were loaded onto a trap column (PepMap Acclaim C18, 5 mm × 300 μm ID, 5 μm particle size, 100 Å pore size, Thermo Scientific) at a flow rate of 25 μl/min using 0.1% TFA as mobile phase.

After 5 minutes, the trap column was switched in line with the analytical column (PepMap Acclaim C18, 500 mm × 75 μm ID, 2 μm particles, 100 Å, Thermo Scientific operated at 30 °C or Aurora Ultimate C18 25cm × 75 μm ID, 1.7 μm particles, 120 Å, with integrated emitter, Ionopticks operated at 50 °C). Peptides were eluted using a flow rate of 230 or 300 nl/min, starting with the mobile phases 98% A (0.1% formic acid in water) and 2% B (80% acetonitrile, 0.1% formic acid) and linearly increasing to 35% B over the next 60 or 120 minutes respectively, followed by an increase to 95% B in 1.7 min, a 4-min hold at 95% B, and re-equilibration with 2% B for three column volumes (equilibration factor of 3.0).

The Orbitrap Exploris 480 mass spectrometer was operated in data-dependent mode, performing a full scan (m/z range 350-1200, resolution 60,000, norm. AGC target 300%) at 3 different compensation voltages (CV −45, −60, −75), followed each by MS/MS scans of the most abundant ions for a cycle time of 0.9 seconds per CV. MS/MS spectra were acquired using HCD collision energy of 30, isolation width of 1.2 m/z, orbitrap resolution of 30,000, norm. AGC target 200%, minimum intensity of 25,000 and maximum injection time of 100 ms. Precursor ions selected for fragmentation (include charge state 2-6) were excluded for 45 s. The monoisotopic precursor selection filter and exclude isotopes feature were enabled.

The Orbitrap Astral was operated in data-independent mode, performing a full scan in the Orbitrap every 0.6 s (m/z range 380–980 m/z; resolution 240,000; AGC target 1,000,000, maximum injection time 5 ms). MS/MS spectra were acquired in the Astral analyser by isolating 4 Da windows across 380–880 m/z (resulting in 124 scan events per cycle) and fragmenting precursor ions with normalized HCD collision energy of 25% with a maximum injection time of 5 ms or until an AGC target of 30,000 was reached. Fragment ions ranging from 150–2000 m/z were acquired.

### Data Processing MS protocol

#### Exploris

For peptide identification, the RAW-files were loaded into Proteome Discoverer (version 2.5.0.400, Thermo Scientific). All MS/MS spectra were searched using MSAmanda version 2.0.0.19924 (Dorfer V. et al., J. Proteome Res. 2014 Aug 1;13(8):3679-84). The peptide and fragment mass tolerance was set to ±10 ppm, the maximum number of missed cleavages was set to 2, using tryptic enzymatic specificity without proline restriction. The RAW-files were searched against the *C.reinhardtii* v6.1 Phytozome database (30,357 sequences; 24,276,980 residues), supplemented with common contaminants using the following modifications: Carbamidomethylation of cysteine was set as fixed modification and oxidation of methionine, deamidation of asparagine and glutamine, phosphorylation on serine, threonine and tyrosine, ubiquitylation residue on lysine, glutamine to pyro-glutamate conversion at peptide N-terminal glutamine and acetylation on the protein N-terminus were set as variable modifications. The localization of the post-translational modification sites within the peptides was performed with the tool ptmRS, based on the tool phosphoRS^65^.

The result was filtered to 1 % FDR on PSM and protein level using the Percolator algorithm^66^ integrated in Proteome Discoverer. Additionally, an Amanda score cut-off of at least 150 was applied. Proteins were filtered to be identified by a minimum of 2 PSMs in at least 1 sample. Protein areas were computed in IMP-apQuant^67^ by summing up unique and razor peptides. Resulting protein areas were normalized using iBAQ^68^. Match-between-runs (MBR) was applied for peptides with high confident peak area that were identified by MS/MS spectra in at least one run. Proteins were filtered to be identified by a minimum of 3 quantified peptides.

#### Astral

Astral DIA data were analyzed in Spectronaut 20.1^68^ (Biognosys). Trypsin/P was specified as a proteolytic enzyme, and up to 2 missed cleavages were allowed in the Pulsar directDIA+ search. Dynamic mass tolerance was applied for calibration and the main search. The RAW files were searched against the *C. reinhardtii* v6.1 Phytozome database (30,357 sequences; 24,276,980 residues), supplemented with common contaminants using the following modifications: Carbamidomethylation of cysteine was searched as fixed modification, whereas oxidation of methionine and acetylation at protein N-termini were defined as variable modifications. Peptides with a length between 7 and 52 amino acids were considered, and results were filtered using Spectronaut default filtering criteria (Precursor Qvalue<0.01, Precursor PEP<0.2, Protein Qvalue <0.01 per Experiment and <0.05 per Run, Protein PEP<0.75). Quantification was performed as specified in Biognosys BGS Factory Default settings, grouping peptides by stripped sequence and performing protein inference using IDPicker. Cross-run normalization in Spectronaut was deactivated due to subsequent mode normalization.

Spectronaut results were exported using Pivot reports on the protein and peptide level and converted to Microsoft Excel files using our in-house software MS2Go (https://ms.imp.ac.at/?action=ms2go). For DIA data MS2Go utilizes the Python library msReport (developed at the Max Perutz Labs Proteomics Facility) for data processing. Abundances were normalized by the mode of protein ratios in msReport and missing values were imputed with values obtained from a log-normal distribution with a mean of 100. To compensate for different protein lengths, protein quantification was then normalized using iBAQ^68^. Statistical significance of differentially expressed proteins was determined using limma^69^.

The mass spectrometry proteomics data have been deposited to the ProteomeXchange Consortium via the PRIDE^70^ partner repository with the dataset identifier PXD072529.

### VIA1 and VIPP1 structure modeling

CrVIA1 signal peptide sequence was predicted using TargetP 2.0^28^ and Wolf PSORT^29^, while its transmembrane domains were predicted using DeepTMHMM^36^. The predicted tertiary protein structure of VIA1 and VIPP1, as well as the interactions between the two proteins, was modeled by AlphaFold3 (AF3)^71^ using the protein sequences lacking their predicted chloroplast signal peptide. Graphical modifications on the protein structures were done using ChimeraX-1.4^72^.

AF3 structural models of VIA1-VIPP1 were used as input for Pickluster^42^ and Alphabridge^41^ analyses. Pickluster was employed to identify recurrent interaction patches across the top-ranked AlphaFold3 models, using default parameters, and visualized with ChimeraX-1.4. Alphabridge was used to detect inter-residue contacts stabilized by hydrogen bonds and salt bridges between the two proteins, with default thresholds for contact distance and bond geometry. Interchain contacts identified by Alphabridge were visualized as chord diagrams representing connections between the corresponding interface regions.

### Molecular Cloning (*C. reinhardtii*)

The plasmid pRAM315 was generated via In-Fusion cloning (Takara) using pRAM254 as the backbone; pRAM254 contained the VIA1 promoter (1,500 bp upstream of the 5′UTR), the 5′UTR, the full VIA1 coding sequence (CDS), a NeonGreen sequence, and a 3×FLAG epitope in frame to the C-terminus of VIA CDS. Two PCRs were performed to amplify the genomic locus of VIA1 from genomic DNA, with the primers oRS41-42 (TM 65 °C, extension time 4’) and oRS43-44 (TM 63 °C, extension time 3’ 30’’). To clone pRAM315, oRS401-402 were used to amplify the 3XFLAG epitope, oRS403-404 to amplify the 3’UTR from genomic DNA (TM 60 °C, extension time 1’), and oRS405-406 were used to linearize the backbone (pRAM254) and remove the NeonGreen sequence. All remaining plasmids were assembled via Golden Gate cloning, following the strategy and toolkit developed in^73^. Briefly, pCSB192 was used as the backbone to insert the coding sequences of *mutVIA1*, *SynVIA1*, and *AtVIA1*, generating level-0 constructs. In the subsequent reaction, the 16S promoter and RbcL terminator/3′UTR were added using plasmids pStA0_16S and pStA0_TrbcL, respectively, with pGT402 as the backbone, yielding level-1 plasmids. In the final step, right and left homology arms targeting the *psbH* locus were incorporated from pHT150 and pHT146, respectively, along with the spectinomycin resistance cassette (*aadA*) from pCSB176. The final level-2 and final constructs were assembled using pCSB192 as the backbone. All plasmids were sequence-verified prior to transformation into Chlamydomonas. A complete list of plasmids and primers used in this study can be found in Tables S3 and S5, respectively.

### Molecular Cloning (*Synechocystis sp.* pcc 6803)

A plasmid containing the *SynVIA1* gene fused to a C-terminal triple FLAG tag and marked with a chloramphenicol resistance cassette (Cmᴿ) was assembled from five PCR-amplified fragments. The downstream fragment was amplified from Synechocystis genomic DNA using primer pair 446/447. The *SynVIA1* gene with 1000 bp of its upstream region was amplified using primers 442/443 and Synechocystis genomic DNA as the template. The plasmid backbone containing the ColE1 origin of replication was amplified from pJET1.2 using primers 448/441. The Cmᴿ cassette was amplified from pACYC184 (New England Biolab) using primers 444/445. All fragments were assembled using the NEBuilder® HiFi DNA Assembly Master Mix.

The SynVIA1 deletion mutant was generated by replacing the *SynVIA1* coding region with a Kanamycin resistance cassette (Kanᴿ) amplified from pUC57-Kanᴿ (GenScript). The upstream and downstream flanking sequences were amplified from Synechocystis genomic DNA using primer pairs 442/449 and 446/447, respectively. The Kanᴿ cassette was amplified using primers 450/445, and the backbone fragment containing the ColE1 origin of replication was amplified from pJET1.2 (REF) with primers 448/441. The four fragments were assembled using the NEBuilder® HiFi DNA Assembly Master Mix. All plasmids were sequence-verified prior to transformation into Synechocystis. Complete deletion of *SynVIA1* was confirmed by PCR using primers 450/445, restriction enzyme digestion with NheI, and sequencing.

### Nuclear transformation (*C. reinhardtii*)

Nuclear transformation was performed by electroporation as described in ref. 18. Briefly, *via1-1* cells were grown in TAP medium to mid-log phase (2–4 × 10⁶ cells mL⁻¹), harvested by centrifugation (800 x g for 5’), and washed twice with 5 mL CHES buffer (10 mM N-cyclohexyl-2-aminoethanesulfonic acid (CHES), pH 9.25, 40 mM sucrose, 10 mM sorbitol). Cells were resuspended in CHES buffer to a final density of 8 × 10⁸ cells mL⁻¹.

For each transformation, 1 µg linearized pRAM315 was mixed with 0.5 µL salmon sperm DNA (10 mg mL⁻¹; Invitrogen 15-632-011) and added to 125 µL of washed cells. The mixture was transferred to 2-mm gap electroporation cuvettes and electroporated using a NEPA21 Type II square-pulse electroporator (ref. 19). The electroporation program consisted of two poring pulses (250 V and 150 V, 8 ms each), followed by five transfer pulses (50 ms each) starting at 20 V with a 40% decay (20, 12, 7.2, 4.3, and 2.6 V).

Immediately after electroporation, cells were recovered in 8 mL TAP supplemented with 40 mM sucrose and incubated overnight at room temperature under low light with gentle shaking. Cells were then harvested (800 x g for 5’), resuspended in 1 mL TAP, and plated on TAP agar containing 20 µg mL⁻¹ hygromycin. Hygromycin-resistant colonies were screened for transgene expression by anti-FLAG immunoblotting.

### Chloroplast transformation (*C. reinhardtii*)

Chloroplast biolistic transformation of the *via1-1* strain was performed using a helium-driven particle gun based on the designs described in^74,75^, and following the protocol in^75^. Cells were grown to ∼4 × 10⁶ cells mL⁻¹, harvested by centrifugation (800 x g for 5’), and resuspended in TAP to a final density of 1 × 10⁸ cells mL⁻¹. 300 µL of concentrated cells was spread onto TAP agar plates containing 100 µg mL⁻¹ spectinomycin (Sigma-Aldrich) and air-dried in a laminar-flow hood for ∼15 min. Plates were bombarded with 550-nm gold particles (Seashell Technology) coated with 1 µg plasmid DNA (pRAM321, pRAM537, or pRAM324, Table S3). After bombardment, plates were incubated under continuous light (80 µmol photons m⁻² s⁻¹) for ∼10 days until colonies appeared. Transformants were restreaked at least four times on selective medium to achieve homoplasmy, and transgene expression was verified by RT–qPCR.

### Transformation (*Synechocystis sp.* pcc 6803)

Wild-type *Synechocystis sp.* pcc 6803 grown in BG11 liquid medium supplemented with 6 µg/ml Ferric ammonium citrate and 5mM glucose under continuous white light (∼8 µmol photons m^−2^ sec^−1^). Once saturated, cells were diluted 10x the day before nuclear transformation. On the day of transformation, cells were harvested when the culture reached an optical density at 730 nm of 0.2–0.6. For each transformation, 5 mL of culture was collected by centrifugation at 3000 x g for 5 minutes. The cell pellet was resuspended in 1 mL of BG11 medium supplemented with iron and transferred to a microcentrifuge tube. Cells were centrifuged again at 2000 x g for 2 minutes and resuspended in 1 mL of BG11 medium. Aliquots of 100 μL were prepared for each transformation. For each transformation, at least 200 ng of plasmid DNA was added to the cell suspension. The transformation mixtures were incubated in the dark for 5 hours. Following incubation, cells were plated on selective BG11 agar plates containing a limiting concentration of the appropriate antibiotic. Colonies were typically observed after 3–4 weeks.

## Data availability

Protein sequences, alignments, phylogenies, and datasets used for phylogenetic analyses have been deposited to Figshare at https://figshare.com/s/c411a207630c1d30d379.

The full phylogeny with species distribution is available via the Interactive Tree of Life (iTOL) at https://itol.embl.de/shared/1WjO4fmjYiy3z.

The mass spectrometry proteomics data have been deposited to the ProteomeXchange Consortium via the PRIDE partner repository^70^ with the dataset identifier PXD072529.

## Supporting Information

**Figure S1.**
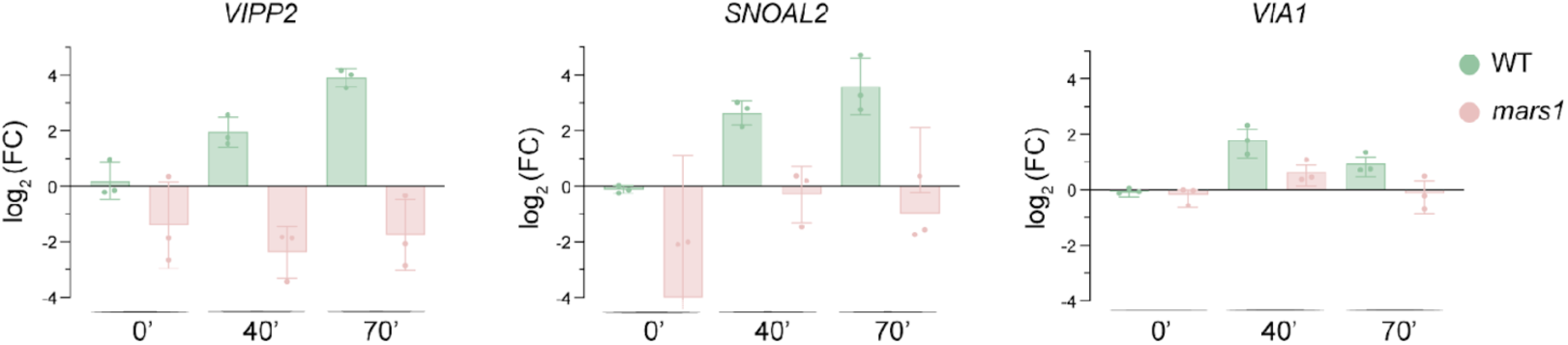
*VIA1* gene expression is upregulated upon cpUPR activation in a MARS1-dependent manner. WT and *mars1* knockdown cells were exposed to high light (>1000 μE) for 0, 40, and 70 minutes. The transcript levels of *VIA1* and two cpUPR marker genes (*VIPP2* and SNOAL) were assessed via RT-qPCR. Expression levels are shown as log₂ fold change relative to WT.

**Figure S2.**
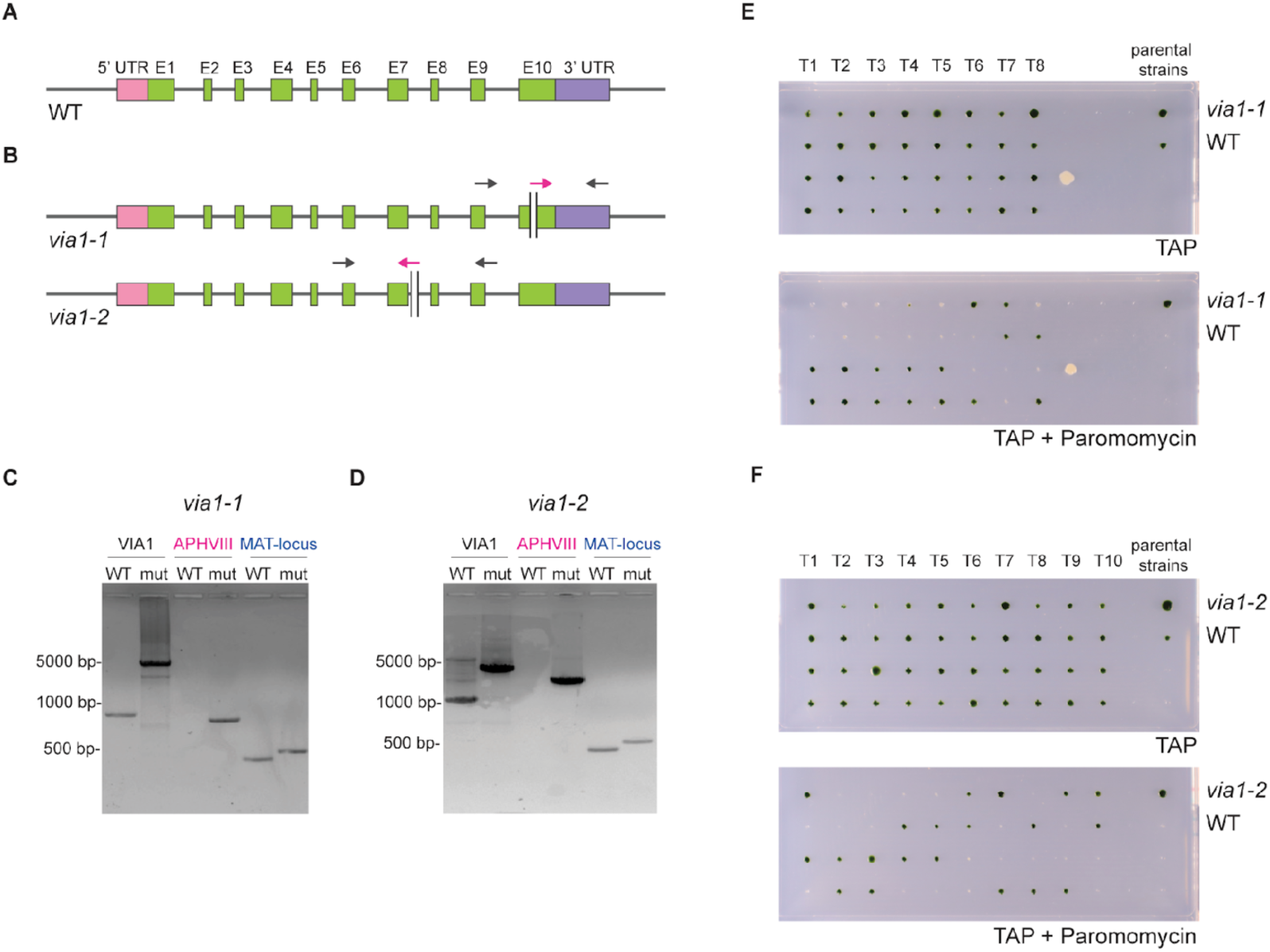
Validation of mutagenic insertion in Chlamydomonas *via1-1* and *via1-2* by PCR and segregation analysis. **(A)** Schematic representation of the *VIA1* transcript: exons are shown in green, 5’UTR in pink, and 3’UTR in purple. **(B)** Diagram illustrating the mutagenic insertion carrying the gene conferring paromomycin resistance (*APHVIII*) in *via-1* and *via1-2*. The mutagenic cassette (represented by black vertical lines) is located in exon 10 and intron 7, respectively. Its orientation is 5’→3’ in *via1-1* and 3’→5’ in *via1-2*. Arrows indicate primers used for the PCRs shown in panels C and D. **(C-D)** PCRs validating the mutagenic insertion in *via1-1* (panel C) and *via1-2* (panel D): The first PCR (*VIA1*) amplifies a region between black arrowheads in the diagrams. The size difference between WT and mutant amplicons reflects the presence of the cassette in mutants. The second PCR (*APHVIII*) amplifies the region between black and pink arrowheads, producing an amplicon only in mutants. The third PCR (*MAT-locus*) is a mating-type-specific PCR used as a control to ensure the quality of the employed genomic DNA. **(E-F)** Segregation analysis of the mutagenic cassette in tetrads (T1–T10) derived from backcrosses of *via1-1* **(E)** or *via1-2* **(F)** to the WT strain. Tetrads were re-arrayed on rectangular TAP agar plates. The upper section of each panel shows the tetrads growing on TAP agar, while the lower section displays the tetrads growing on TAP agar supplemented with paromomycin

**Figure S3.**
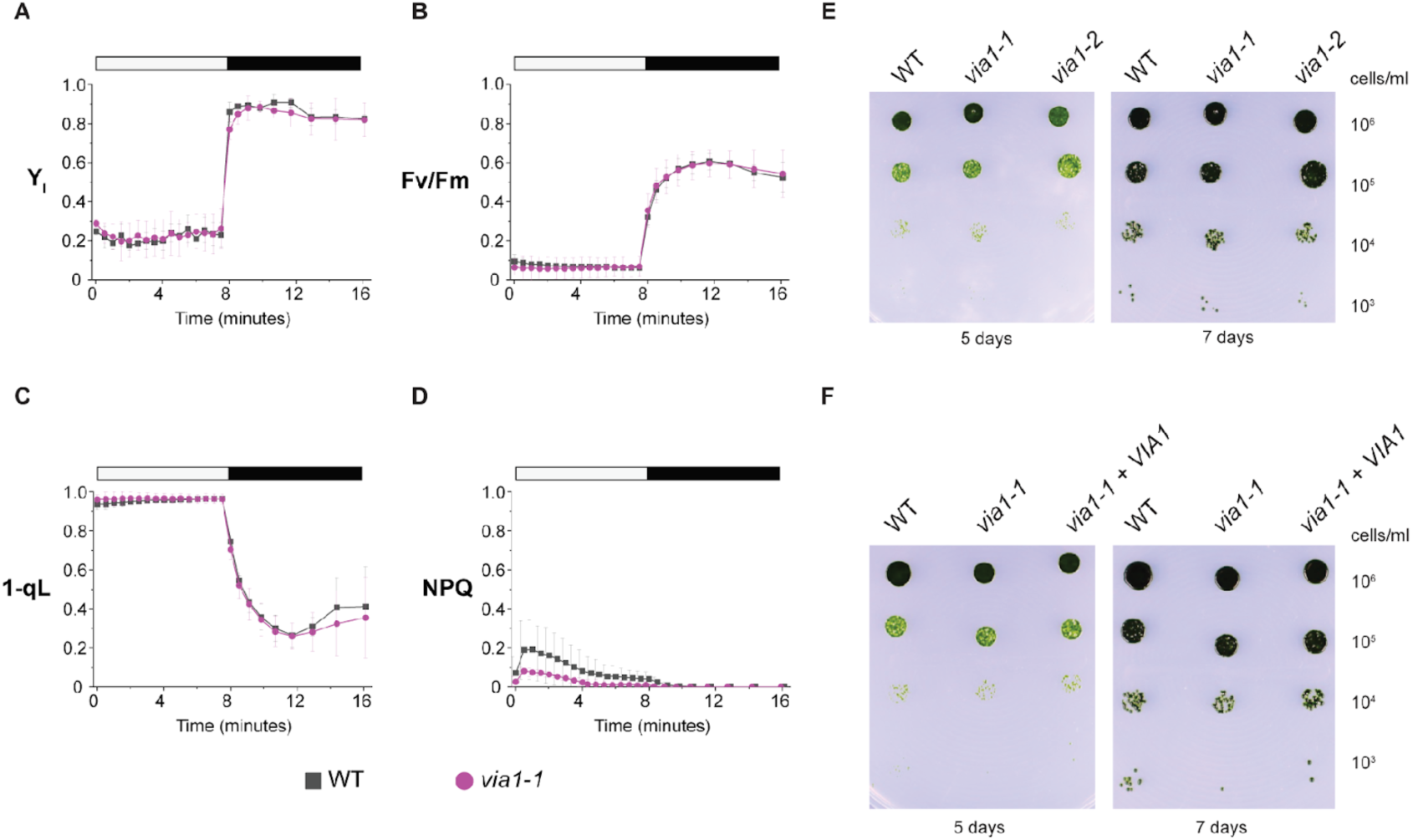
Photosynthetic measurements in Chlamydomonas WT and *via1-1* cells and growth assay in normal light conditions. **(A-B)** Photosystem I quantum yield (YI) **(A)** and Photosystem II quantum yield (Fv/Fm) **(B)** were monitored during an 8-minute exposure to actinic light at an intensity of 1000 µmol photons m⁻² s⁻¹, followed by an 8-minute dark period. **(C-D)** The plastoquinone redox state was assessed by analyzing the fluorescence parameters 1-qL **(C)** and NPQ **(D)**. Data represent means ± standard deviation from four independent measurements. **(E-F)** Growth assay on TAP agar showing the growth of WT, *via1-1*, and *via1-2* **(E)**, and WT, *via1-1*, and *via1-1* + VIA1 **(F)** over 7 days. Liquid cultures were serially diluted to the indicated concentrations, spotted on TAP agar, grown under normal light (∼80 µmol photons m⁻² s⁻¹), and imaged 5- and 7-days post-spotting.

**Figure S4.**
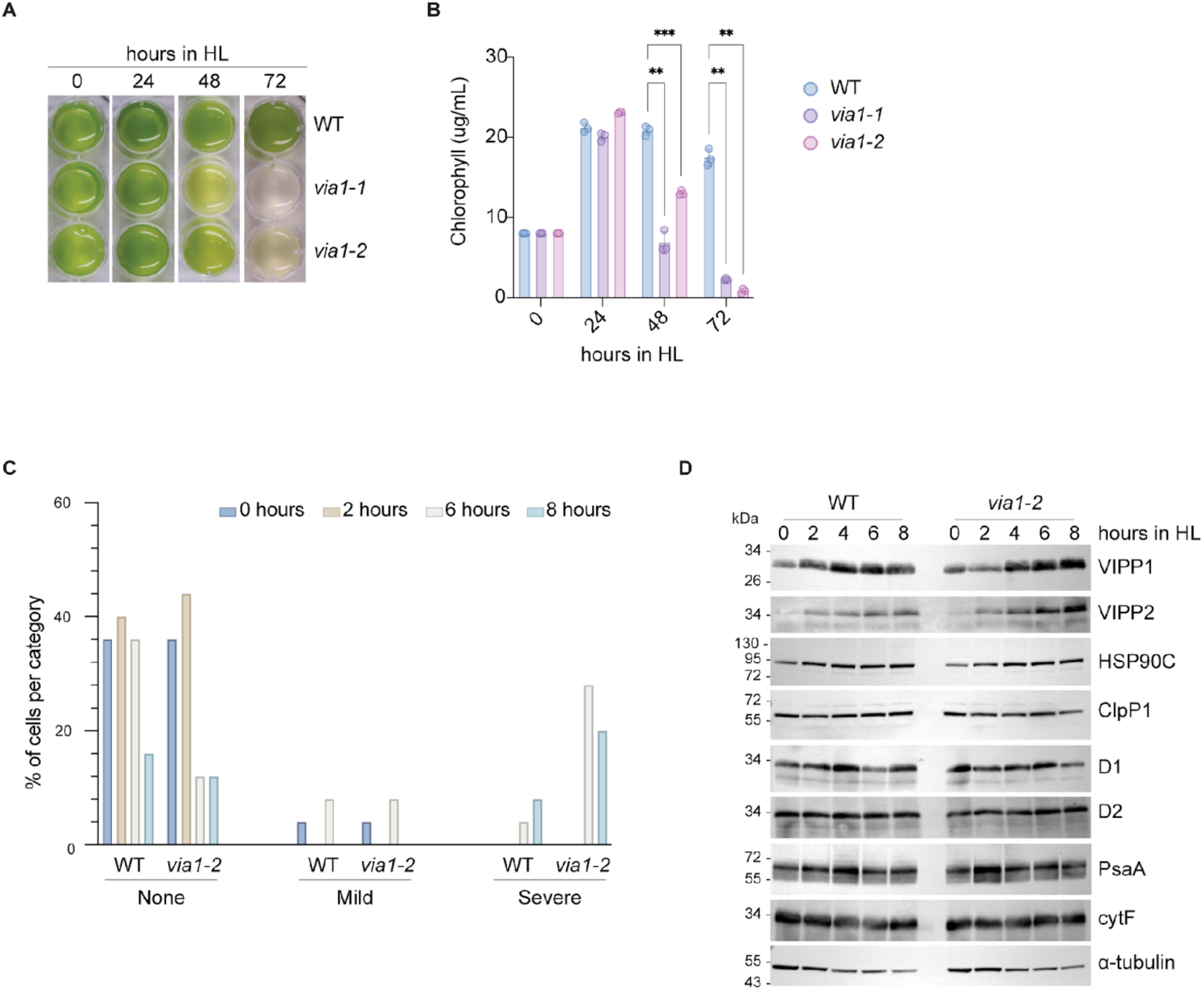
Additional characterization of *via1* mutant strains. **(A)** Images of liquid TAP cultures and **(B)** relative chlorophyll quantification of WT, *via1-1*, and *via1-2* at various time points during a 3-day HL treatment (∼800 µmol photons m^−2^ s^−1^). Three biological replicates were used. **(C)** Distribution of thylakoid swelling severity (none, mild, severe) in WT and *via1-2* cells imaged in Fig. 2E–F at the indicated time points during HL treatment. **(D)** Time-course immunoblot comparing the accumulation of VIPP1, D1, D2, PsaA, cytF, and ClpP1 in WT versus *via1-2* during HL treatment (0–8 h); α-tubulin serves as a loading control.

**Figure S5.**
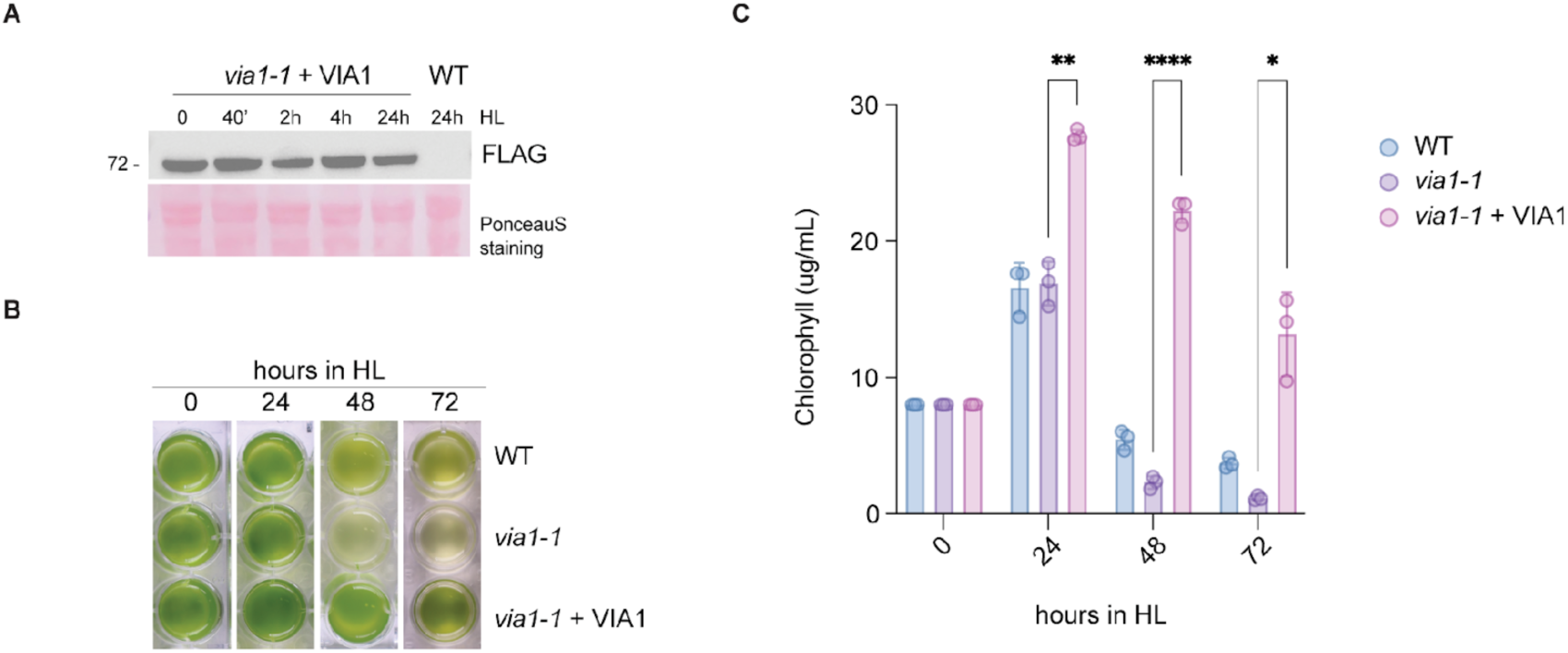
Phenotypic rescue of *via1* upon introduction of VIA1-FLAG transgene. **(A)** FLAG immunoblot analysis of total protein lysates from *via1-1* + *VIA1:FLAG* cells subjected to HL treatment. **(B)** Representative images of liquid TAP cultures of WT, *via1-1*, and *via1-1 + VIA1:FLAG,* and **(C)** relative chlorophyll content measured at the indicated time points over a 3-day HL treatment (∼800 µmol photons m^−2^ s^−1^). Data are from three biological replicates.

**Figure S6.**
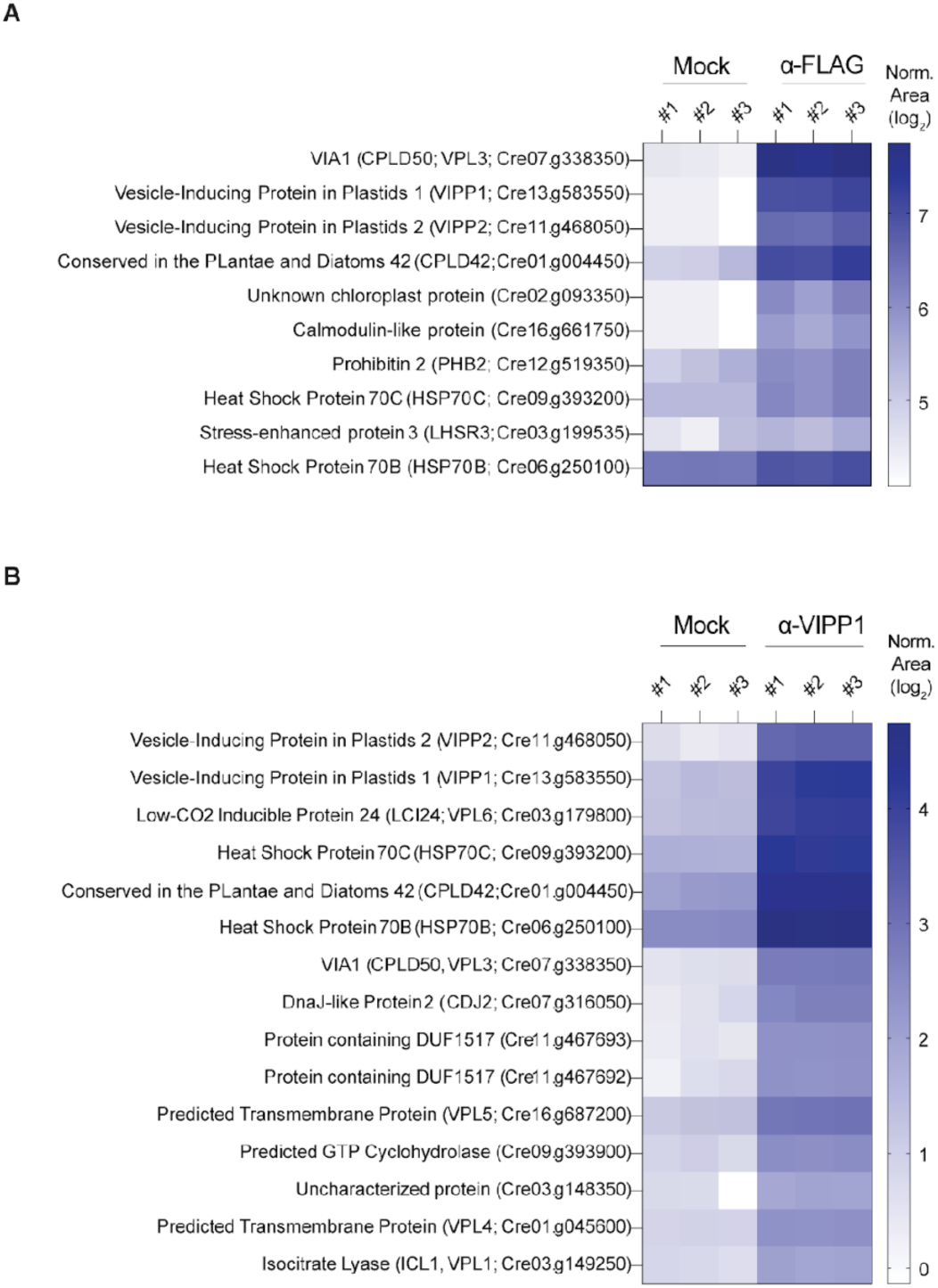
VIA1 interacts with VIPP1 in *Chlamydomonas reinhardtii*. **(A)** Heatmap showcasing proteins reproducibly enriched in FLAG co-immunoprecipitations from *via1* + *VIA1:FLAG* strain (three biological replicates) relative to control samples consisting of WT lysate incubated with FLAG–M2 beads. Heatmap colors indicate the log₂ normalized area (relative abundance), with darker shading denoting higher relative abundance. **(B)** Heatmap of proteins significantly enriched in VIPP1 co-immunoprecipitations from WT cell, shown as log₂ normalized area (relative abundance). Control samples consisted of WT lysates incubated with uncoupled Protein A Dynabeads.

**Figure S7.**
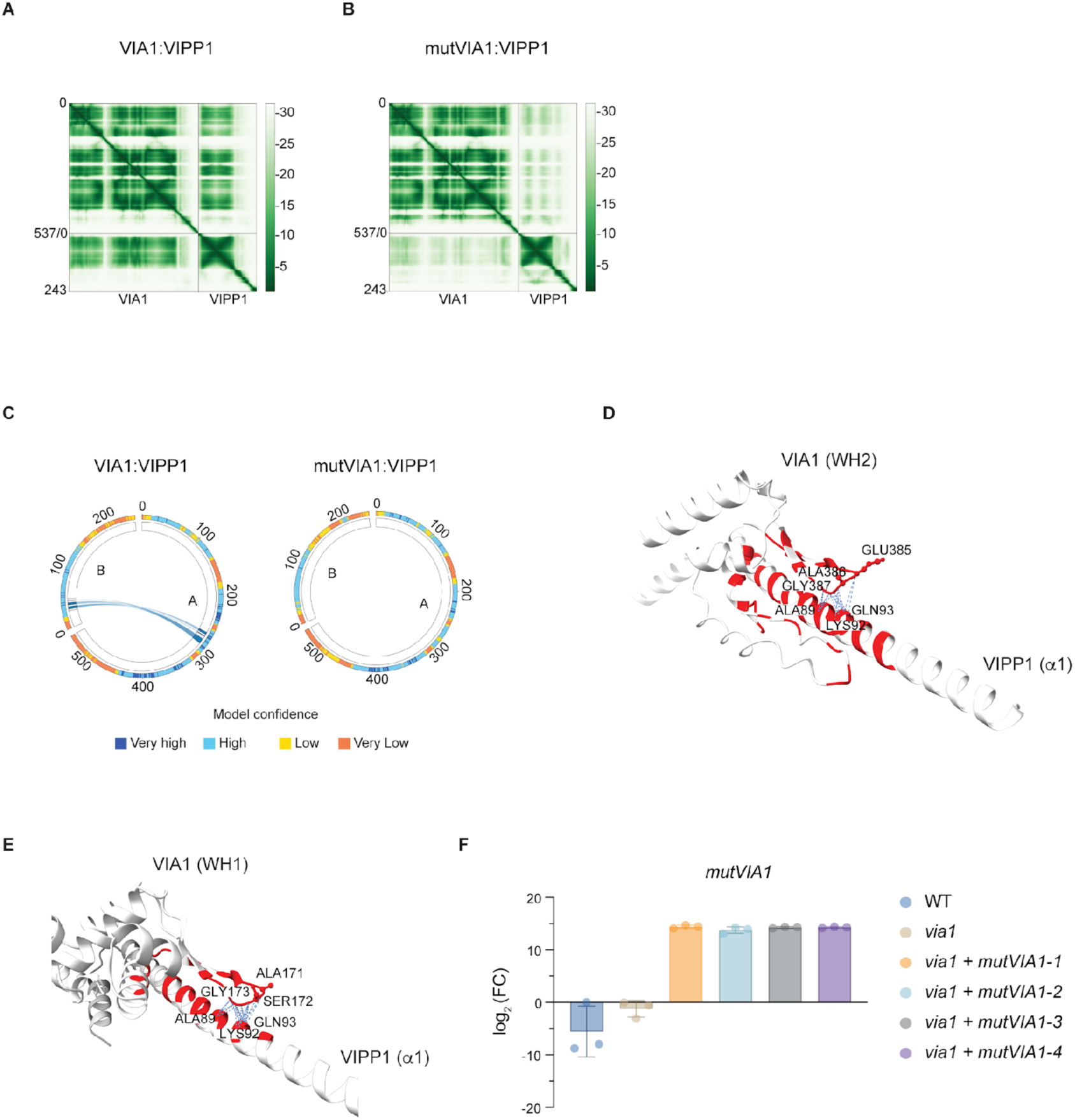
Structural prediction-guided design of VIA1 point mutants targeting the VIA1:VIPP1 interface. **(A-B)** Predicted Aligned Error (PAE) plots from the AlphaFold 3 models of the VIA1–VIPP1 **(A)** and mutVIA1–VIPP1 **(B)** complexes. **(C)** AlphaBridge chord diagrams of predicted residue–residue contacts between VIA1 and VIPP1 and between mutVIA1 and VIPP1. **(D-E)** PickLuster analysis of predicted interaction interfaces between VIPP1 and the WH2 **(D)** or WH1 **(E)** domain of VIA1, with predicted contacts mapped to loop regions targeted for point mutagenesis. **(F)** RT-qPCR analysis of *mutVIA1* transcript levels in WT, *via1*, and four independent *via1* + *mutVIA1* transformant strains. Expression levels are shown as log₂ fold change relative to WT.

**Figure S8.**
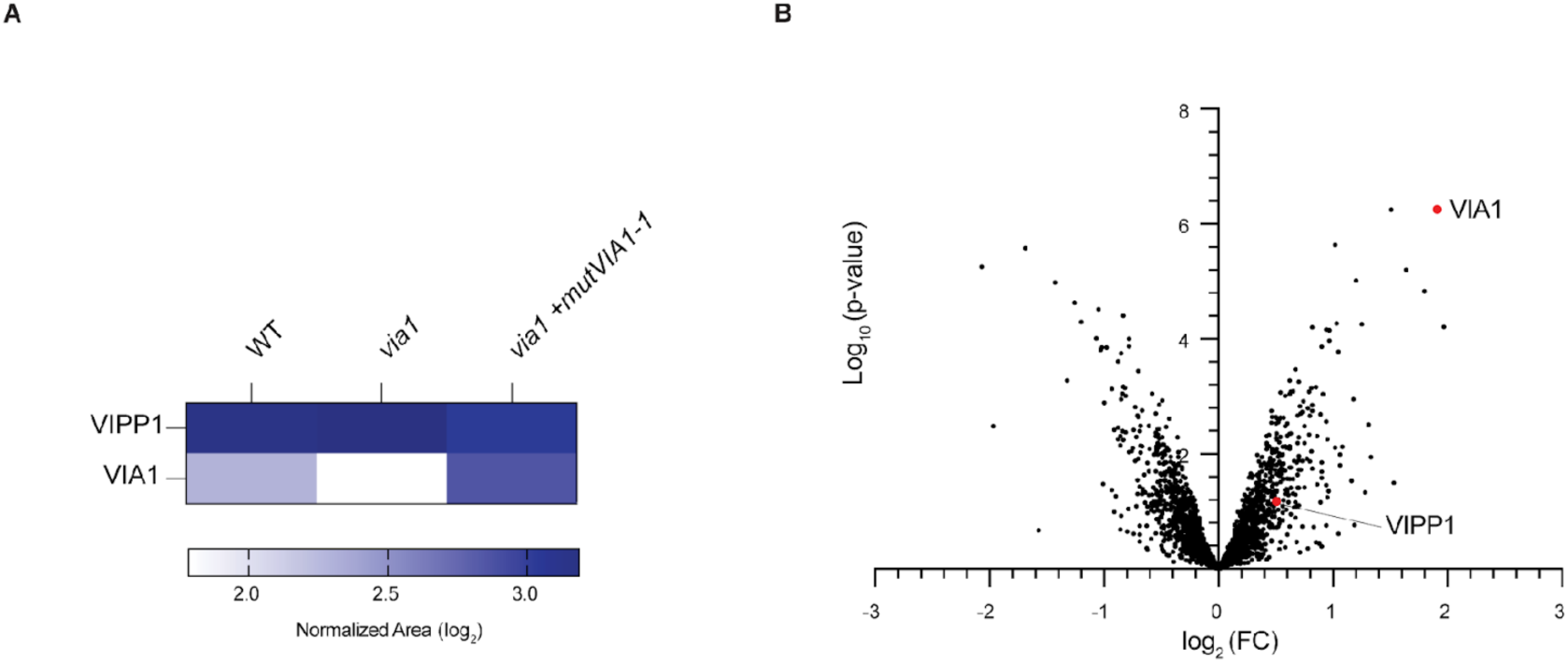
Comparative analysis of VIA1 and VIPP1 levels in VIPP1-co-immunoprecipitation inputs and eluates from WT, *via1*, and *via1* + *mutVIA1* strains. **(A)** Heatmap showing log₂ normalized area (relative abundance) for VIA1 and VIPP1 in the VIPP1 co-immunoprecipitation inputs from WT, *via1*, and *via1* + *mutVIA1* strains. **(B)** Volcano plot of proteins quantified in the WT and *via1* + *mutVIA1* VIPP1 immunoprecipitation datasets. The x-axis shows log₂ fold change, and the y-axis shows −log₁₀(p-value). VIA1 and VIPP1 are highlighted in red.

**Figure S9.**
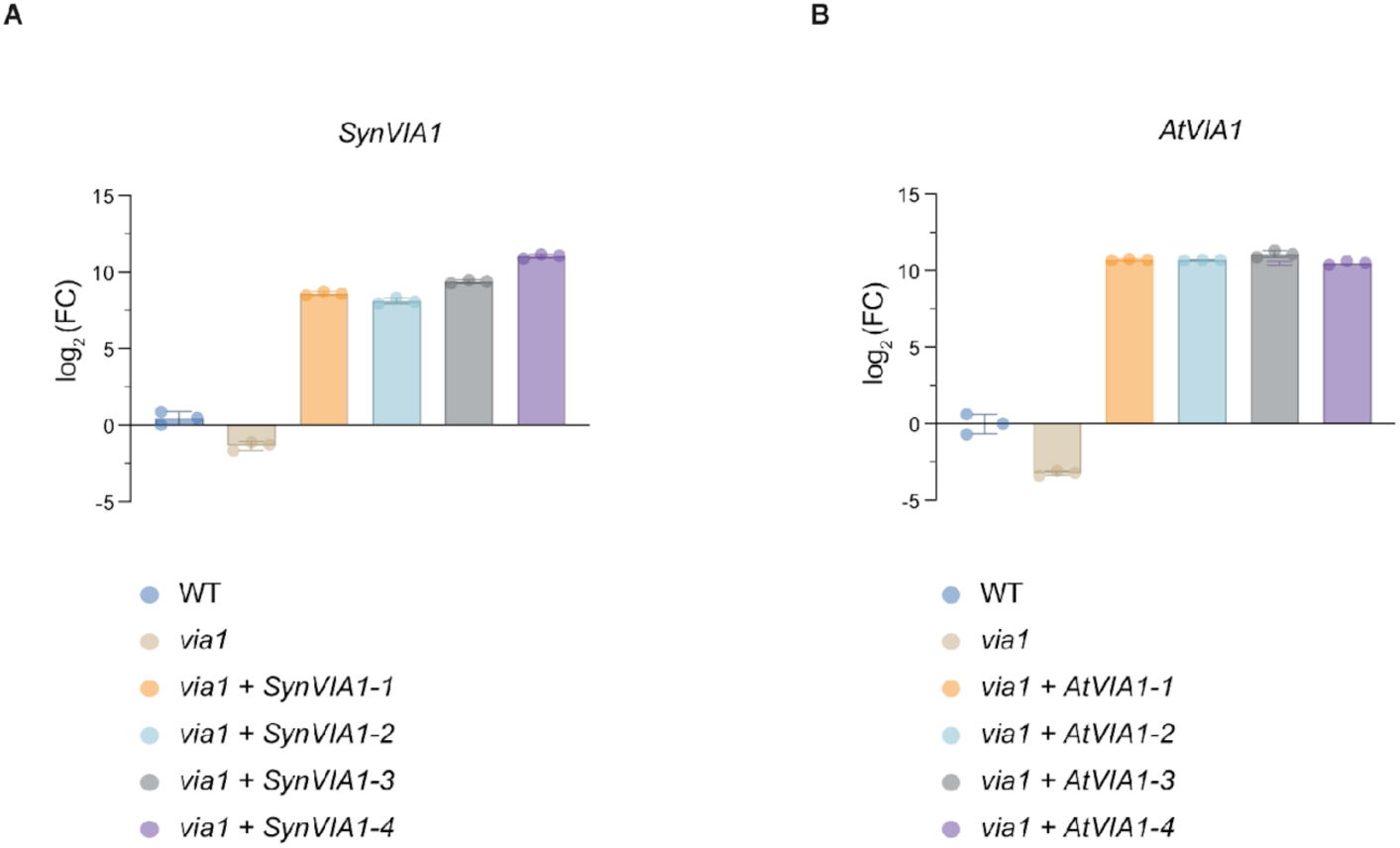
Expression of *VIA1* orthologs in cross-species complementation strains. qRT–PCR analysis of VIA1 ortholog expression from *Synechocystis sp.* PCC 6803 **(A)** and *Arabidopsis thaliana* **(B)** introduced into the Chlamydomonas *via1* mutant via biolistic chloroplast transformation. Transcript levels are shown as log₂ fold change relative to WT. Four independent transformants were analyzed for each ortholog.

**Figure S10.**
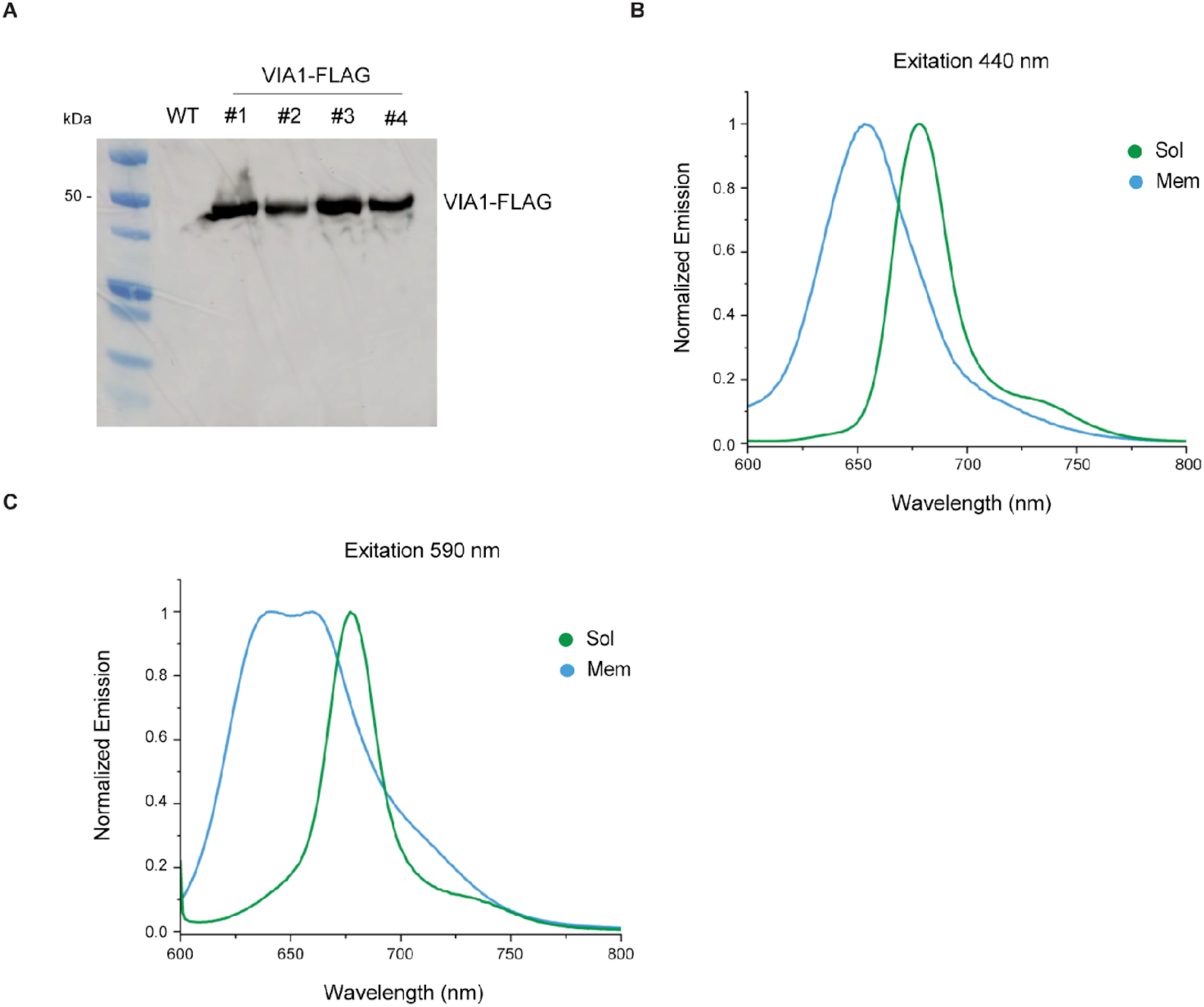
Generation and biochemical characterization of a VIA1-FLAG tagged strain in *Synechocystis sp.* PCC 6803. **(A)** The endogenous *VIA* gene (*slr1603*) was tagged with a C-terminal 3XFlag. Four independent clones obtained after complete segregation of the tagged allele were verified by FLAG immunoblot analysis. Clone #1 was used in the subsequent experiments. **(B)** Normalized fluorescence emission spectra excited at 440 nm (Chl excitation) reveal trace Chl levels in the soluble fraction and confirm the absence of PBS signal in the membrane fraction. **(C)** Normalized fluorescence emission spectra excited at 590 nm (PBS excitation) confirm the absence of PBS in the membrane fraction and reveal only trace Chl levels in the soluble fraction.

**Figure S11.**
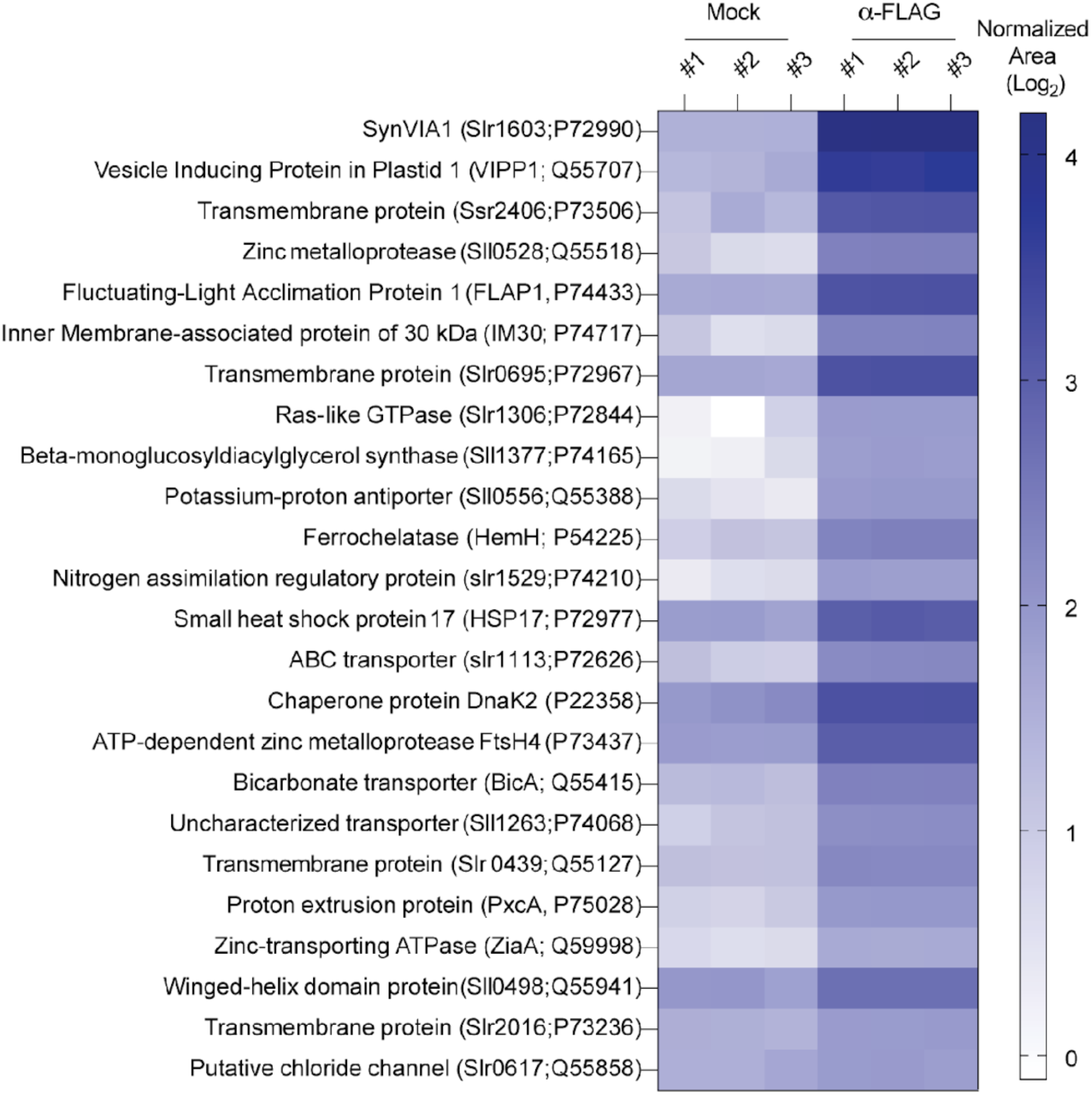
VIA1 interacts with VIPP1 in *Synechocystis sp.* PCC 6803. Heatmap of proteins reproducibly enriched in FLAG co-immunoprecipitations from a *Synechocystis* sp. PCC 6803 strain expressing SynVIA1-FLAG, relative to untagged WT control samples. Heatmap colors represent log₂ normalized area (relative protein abundance), with darker shading indicating higher abundance.

**Figure S12.**
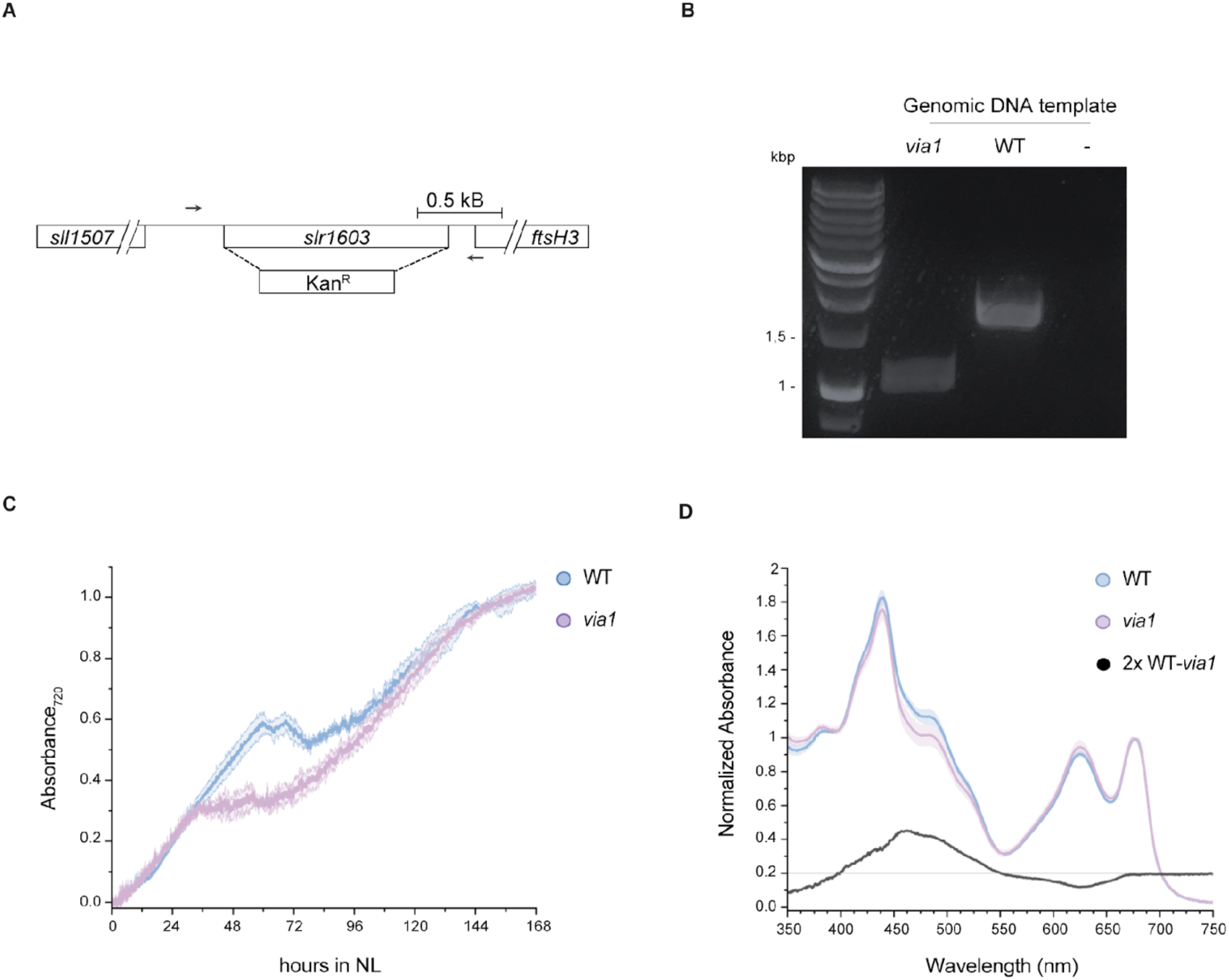
Generation and characterization of a *via1* mutant strain in *Synechocystis sp.* PCC 6803. **(A)** The entire coding sequence of VIA1 (*slr1603*) was replaced by a kanamycin resistance cassette (Kan^R^) to generate an insertional null mutant. Arrows indicate the positions of the genotyping primers used for PCR analysis in panel B (oYM472-473 in Table S5). **(B)** Successful disruption of the VIA1 locus was verified by PCR analysis of the obtained mutant strain. **(C)** Growth analysis of *via1* mutant grown under normal light conditions (NL; 80 µmol photons m⁻² s⁻¹). Data represent the mean optical density at 720 nm (OD_720_) of four independent biological replicates, with shading indicating ± SE. **(D)** Whole-cell absorption spectra of the same cultures. Spectra are normalized to the chlorophyll a absorption peak at 676 nm. The difference spectrum (multiplied by two) between WT and *via1,* which indicates a reduction in cellular carotene levels in *via*1 cells, is shown below as a black line centered at 0.2. Shaded regions indicate standard deviation.

**Figure S13.**
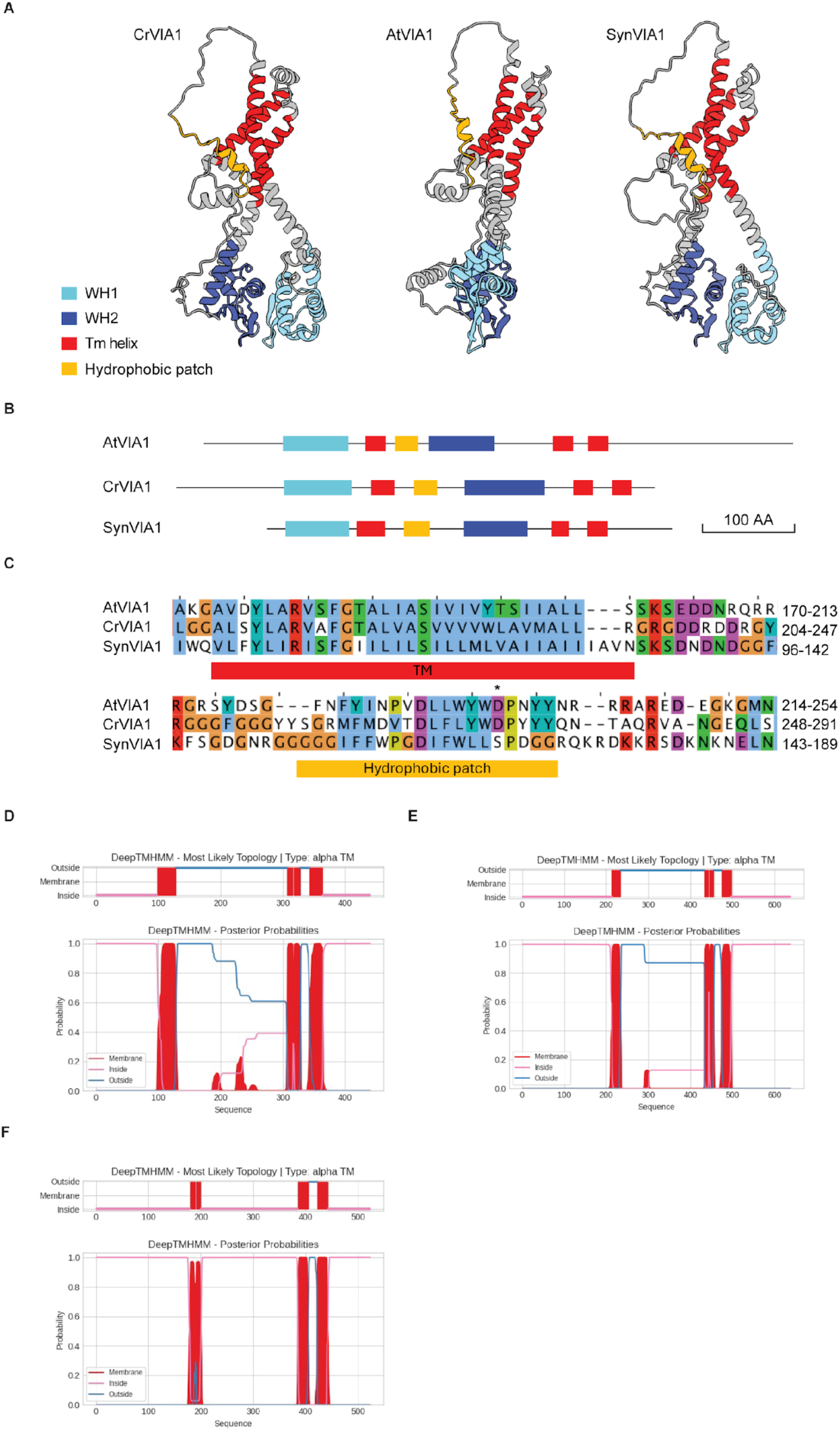
A conserved hydrophobic stretch in VIA1 may form an additional membrane-spanning segment. **(A)** AlphaFold3 models of three VIA1 orthologs showing the two WH domains in close apposition. Domain coloring is consistent across panels B and C. **(B)** The domain organization of AtVIA1, CrVIA1, and SynVIA1 includes a single predicted transmembrane helix between WH1 and WH2, which is inconsistent with both WH domains residing on the same side of the membrane. **(C)** Sequence alignment of the three VIA1 orthologs reveals a conserved hydrophobic stretch immediately following the first transmembrane helix, interrupted by a small number of charged residues. **(D–E)** DeepTMHMM predictions for SynVIA1 **(D)** and CrVIA1 **(E)** assign a non-zero probability to a transmembrane helix in this region. **(F)** The corresponding DeepTMHMM prediction for AtVIA1 assigns zero probability, possibly due to an additional aspartate residue that disrupts the continuity of hydrophobic residues (marked with asterisks in panel C).

**Table S1.**
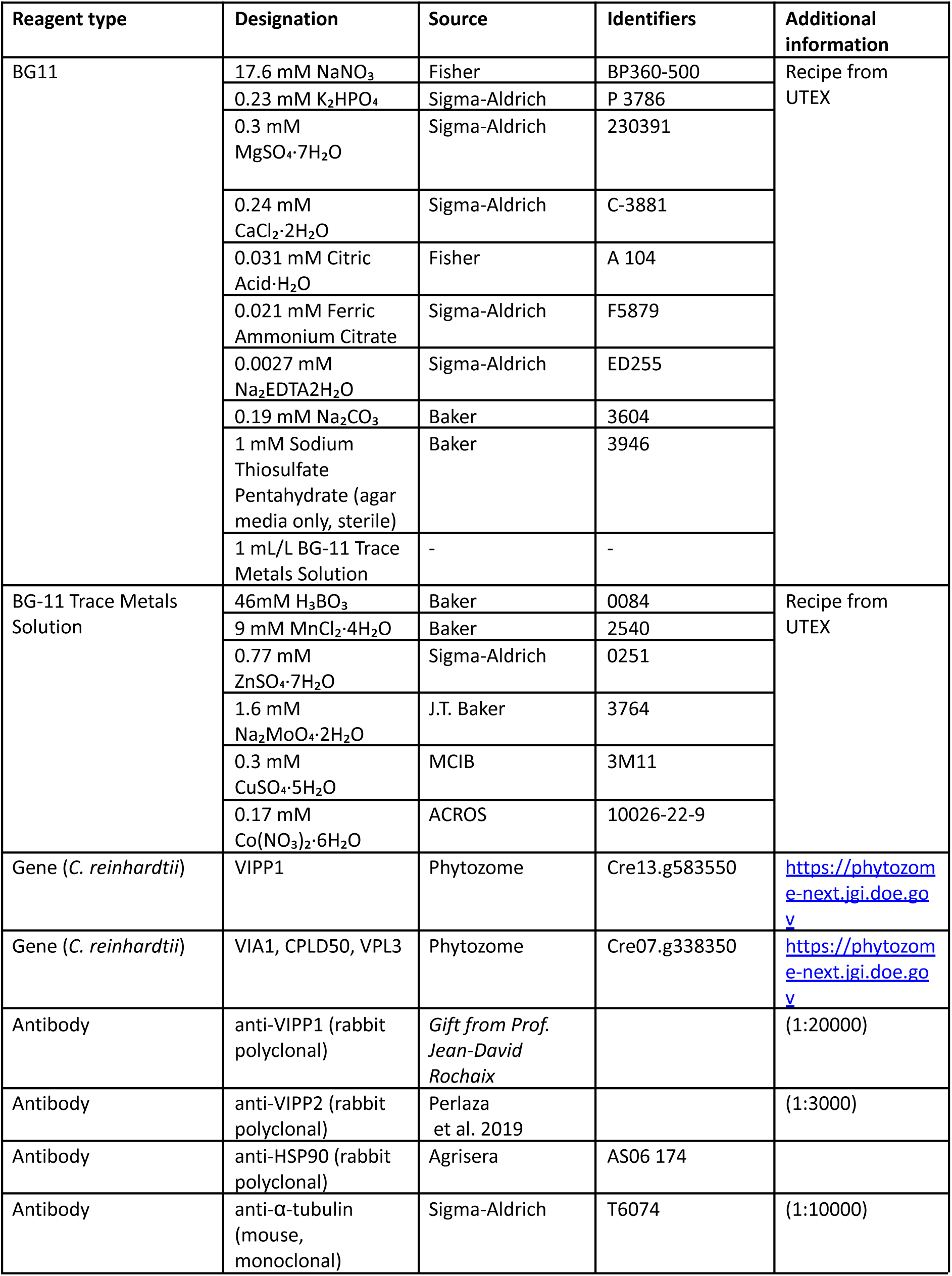
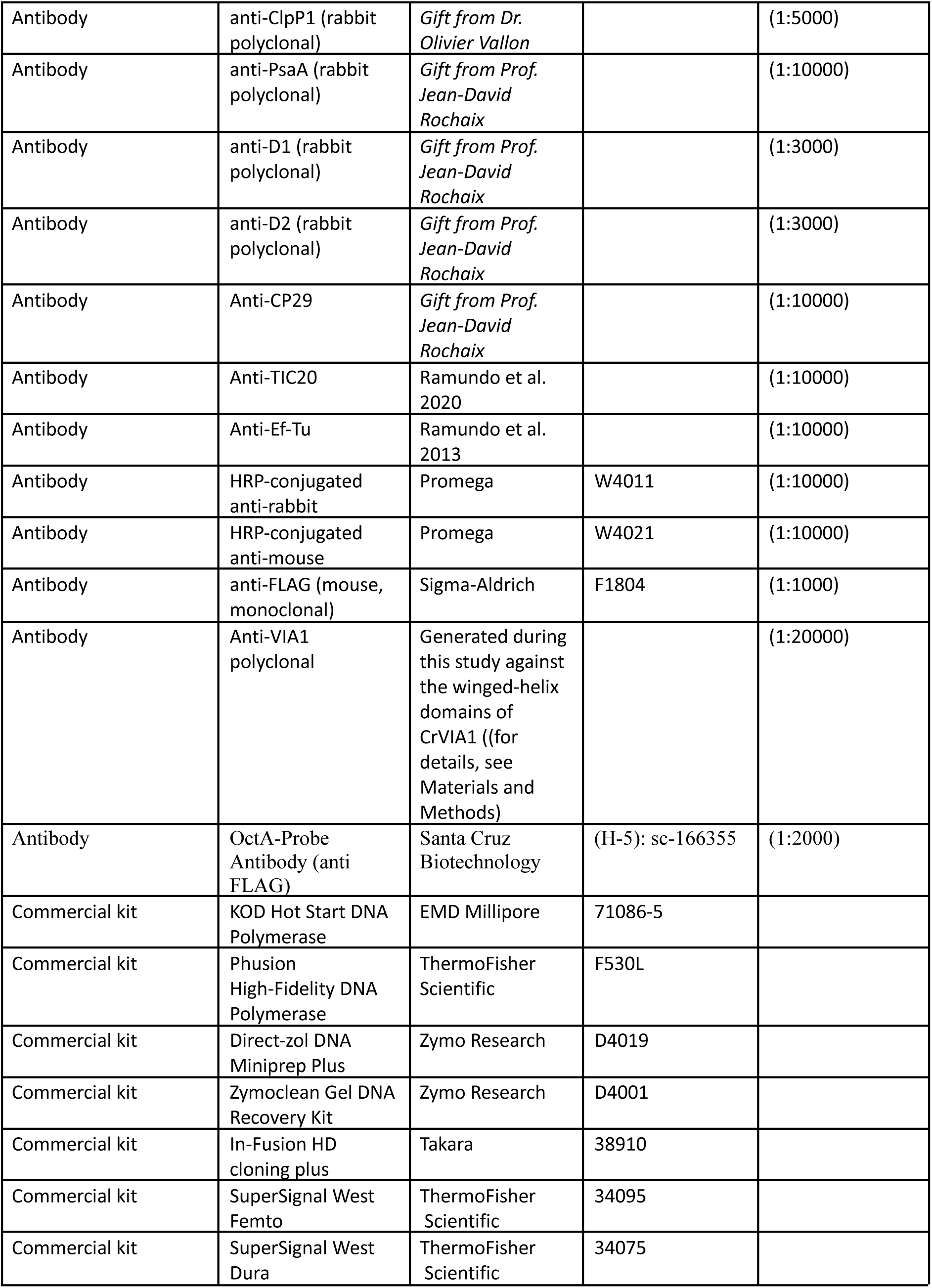
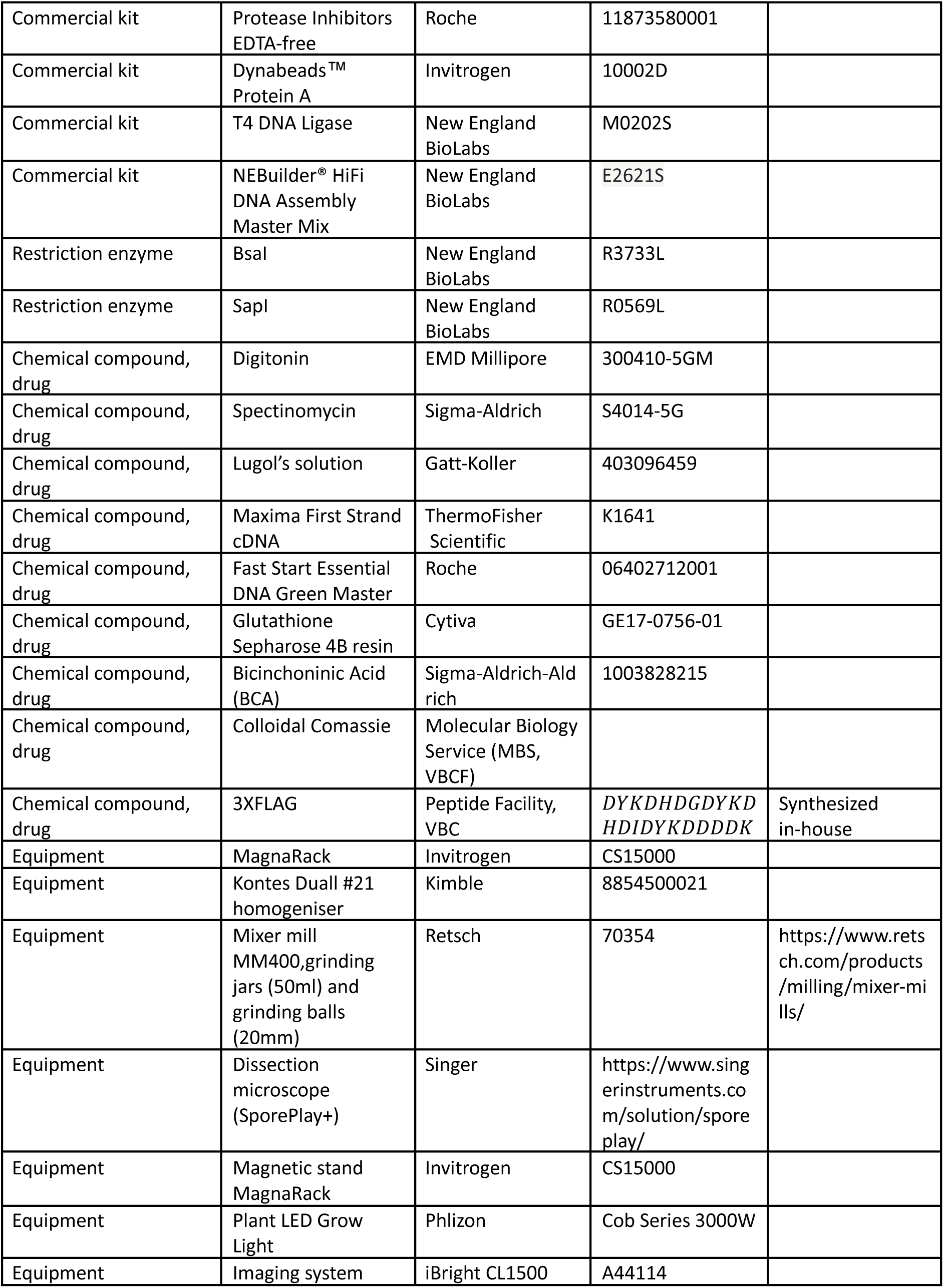
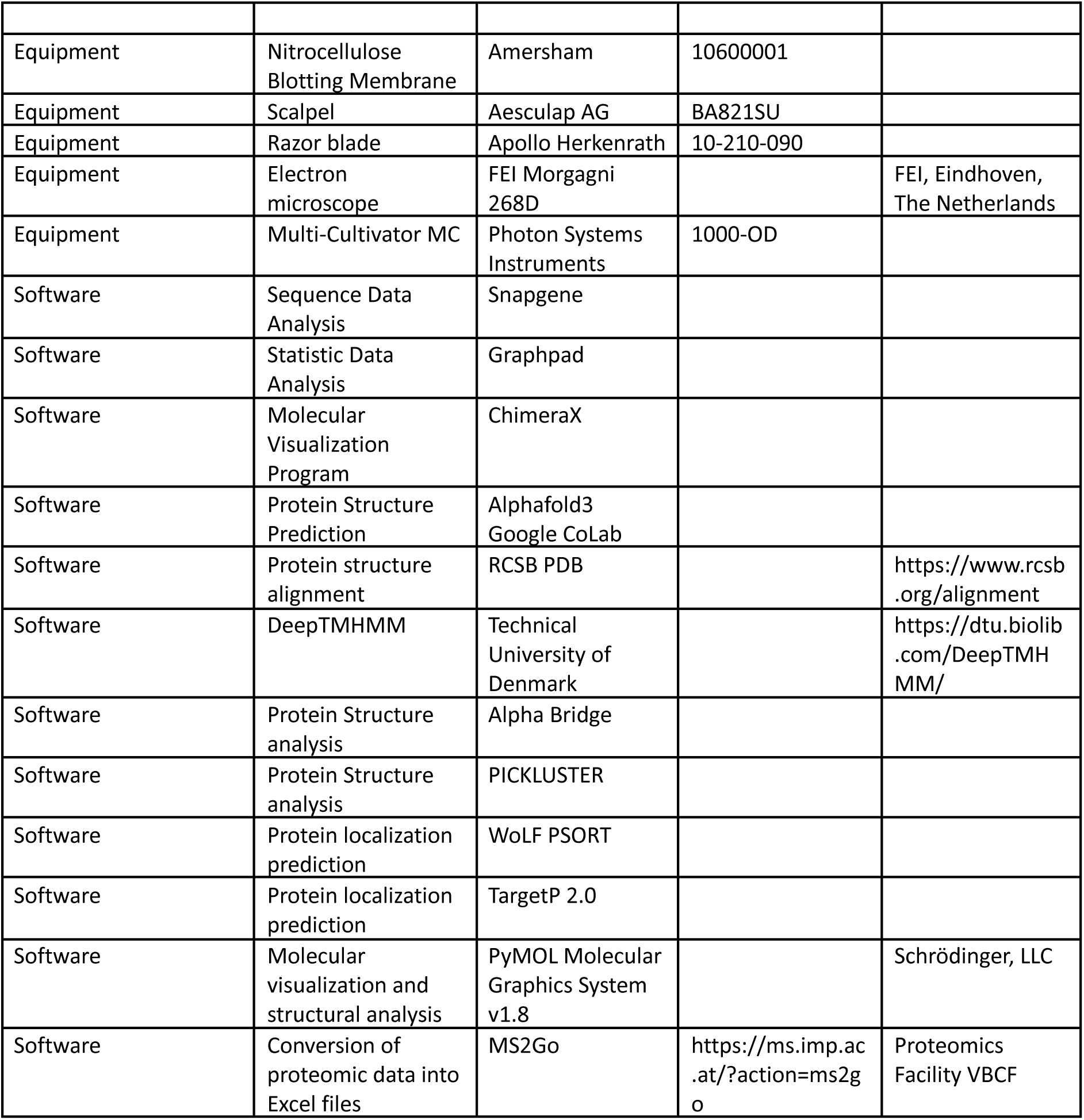
List of reagents used in this study.

**Table S2.** List of all the strains used in this study.

| ID | Strain name | Organism | Description | Comments | Reference |
| --- | --- | --- | --- | --- | --- |
| CrRS166 | CC-5325 | *C. reinhardtii* | wild-type | This strain was employed for all the experiments except for the backcross. |  |
| CrRS197 | XF305 F1 | *C. reinhardtii* | wild-type like progeny from backcross of a CC-5325 strain to a CC-124 strain | This strain was only employed for the backcross experiment. |  |
| CrRS432 | *via1-1* | *C. reinhardtii* | (Available at the Chlamydomonas Resource Center) |  | Cre07.g338350 mutant allele<br>Clip ID: LMJ.RY0402.176599 |
| CrRS433 | *via1-2* | *C. reinhardtii* | (Available at the Chlamydomonas Resource Center) |  | Cre07.g338350 mutant allele<br>Clip ID: LMJ.RY0402.193812 |
| Cr459 | *mars1* | *C. reinhardtii* | Cre16.g692228 (MARS1) mutant |  | Cre16.g692228 mutant allele<br>Clip ID: LMJ.RY0402.195536<br>Perlaza et al. 2019 |
| SPV54.1 | T1 to T8 | *C. reinhardtii* | Plate containing tetrads obtained from backcross of CrRS432 to CrRS197 |  | Strain generated during this study |
| SPV54.2 | T1 to T10 | *C. reinhardtii* | Plate containing tetrads obtained from backcross of CrSR433 to CrRS197 |  | Strain generated during this study |
| Cr528 | *via1-1 + VIA1* | *C. reinhardtii* | *via1-1* expressing VIA1-3XFLAG | Strain employed for the FLAG IP | Strain generated during this study |
| A1 SLM3 | *via1-1 + mutVIA1-1* | *C. reinhardtii* | *via1-1* expressing mutVIA1; mutVIA1 contains the following point mutations A70D S71A G72A | Strain employed for the VIPP1 IP and the HL growth assay | Strain generated during this study |
|  |  |  | E285Q A285D<br>G286V |  |  |
| A3 SLM3 | *via1-1 +*<br>*mutVIA1-2* | *C. reinhardtii* | *via1-1*<br>expressing<br>mutVIA1;<br>mutVIA1<br>contains the<br>following point<br>mutations A70D<br>S71A G72A<br>E285Q A285D<br>G286V | Strain employed<br>for the HL<br>growth assay | Strain generated<br>during this study |
| A3 SLM2 | *via1-1 +*<br>*AtVIA1-1* | *C. reinhardtii* | *via1-1*<br>expressing<br>AtVIA1, the<br>ortholog of VIA1<br>from<br>*Arabidopsis*<br>*thaliana*. | Strain employed<br>for the HL<br>growth assay | Strain generated<br>during this study |
| A11 SLM1 | *via1-1 +*<br>*SynVIA1-1* | *C. reinhardtii* | *via1-1*<br>expressing<br>SynVIA1, the<br>ortholog of VIA1<br>from<br>*Synechocystis sp*<br>*pcc 6803* | Strain employed<br>for the HL<br>growth assay | Strain generated<br>during this study |
| 261 | SynVIA1-FLAG | *E. coli* | DH5a<br>transformed by<br>SynVIA1-FLAG<br>plasmid | Verified by<br>restriction<br>digest and<br>sequencing.<br>CM<sup>R</sup> resistant |  |
| 263 | DSynVIA1 | *E. coli* | DH5a<br>transformed by<br>SynVIA1-FLAG<br>plasmid | Verified by<br>restriction<br>digest and<br>sequencing.<br>CM<sup>R</sup> resistant |  |
| 272 | SynVIA1-FLAG | *Synechocystis*<br>*sp pcc 6803* | Syn transformed<br>with<br>SynVIA1-FLAG<br>plasmid. | Verified by<br>restriction<br>digest. CM<sup>R</sup><br>resistant |  |

**Table S3.** List of all plasmids used in this study.

| Plasmid | Gene | Purpose | Source |
| --- | --- | --- | --- |
| pRAM254 | *pVIA1:VIA1-NeonGreen-3×FLAG* | *via1-1* mutant complementation | Plasmid generated during this study |
| pRAM315 | *pVIA1:VIA1-3×FLAG* | *via1-1* mutant complementation | Plasmid generated during this study |
| pRAM321 | *VIA1_A70D/S71A/G72A/E285/QA285D/G286V* (gene ordered from Twist Bioscience) | *via1-1* mutant complementation | Plasmid generated during this study |
| pRAM324 | *AtVIA1* (gene ordered from Twist Bioscience) | *via1-1* mutant complementation | Plasmid generated during this study |
| pRAM537 | *SynVIA1* (gene ordered from Twist Bioscience) | *via1-1* mutant complementation | Plasmid generated during this study |
| pNS001 | Empty vector | Backbone for cloning pRAM254 and pRAM315 | Gift from Masayuki Onishi |
| pCSB192 | Empty Level 0 and 2 | Cloning of pRAM321, 324, and 537 | Gift from Pawel Mordaka |
| pStA0_16S | *rrn16S* promoter and 5'UTR | Cloning of pRAM321, 324, and 537 | Gift from Pawel Mordaka |
| pStA0_TrbcL | *RbcL* terminator and 3'UTR | Cloning of pRAM321, 324, and 537 | Gift from Pawel Mordaka |
| pGT402 | Empty Level 1 | Cloning of pRAM321, 324, and 537 | Gift from Pawel Mordaka |
| pHT150 | *psbH* right flanking sequence | Cloning of pRAM321, 324, and 537 | Gift from Pawel Mordaka |
| pHT146 | *psbH* left flanking sequence | Cloning of pRAM321, 324, and 537 | Gift from Pawel Mordaka |
| SynVIA1-FLAG | 3x FLAG tag motif with flag linker to C-term of SynVIA1 gene | Co-Immunoprecipitation | Generated by the Mazor lab |
| DSynVIA1 | replacement of *SynVIA1* gene with *Kan<sup>RR</sup>* gene | Growth phenotype | Generated by the Mazor lab |
| pJET1.2 | Plasmid containing *Kan<sup>RR</sup>* gene | Used to clone *Kan<sup>RR</sup>* via PCR | Thermo Scientific |

**Table S4.**
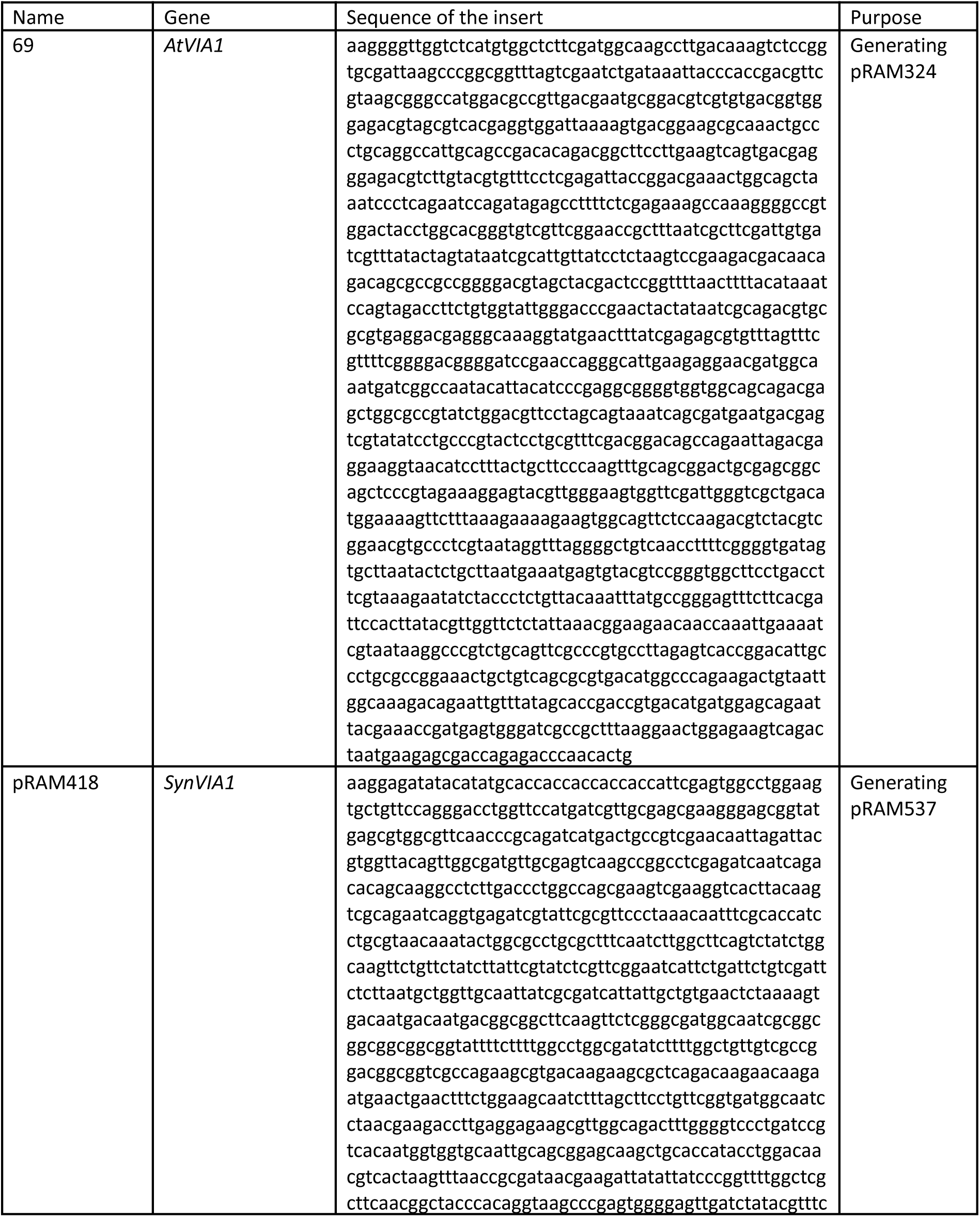
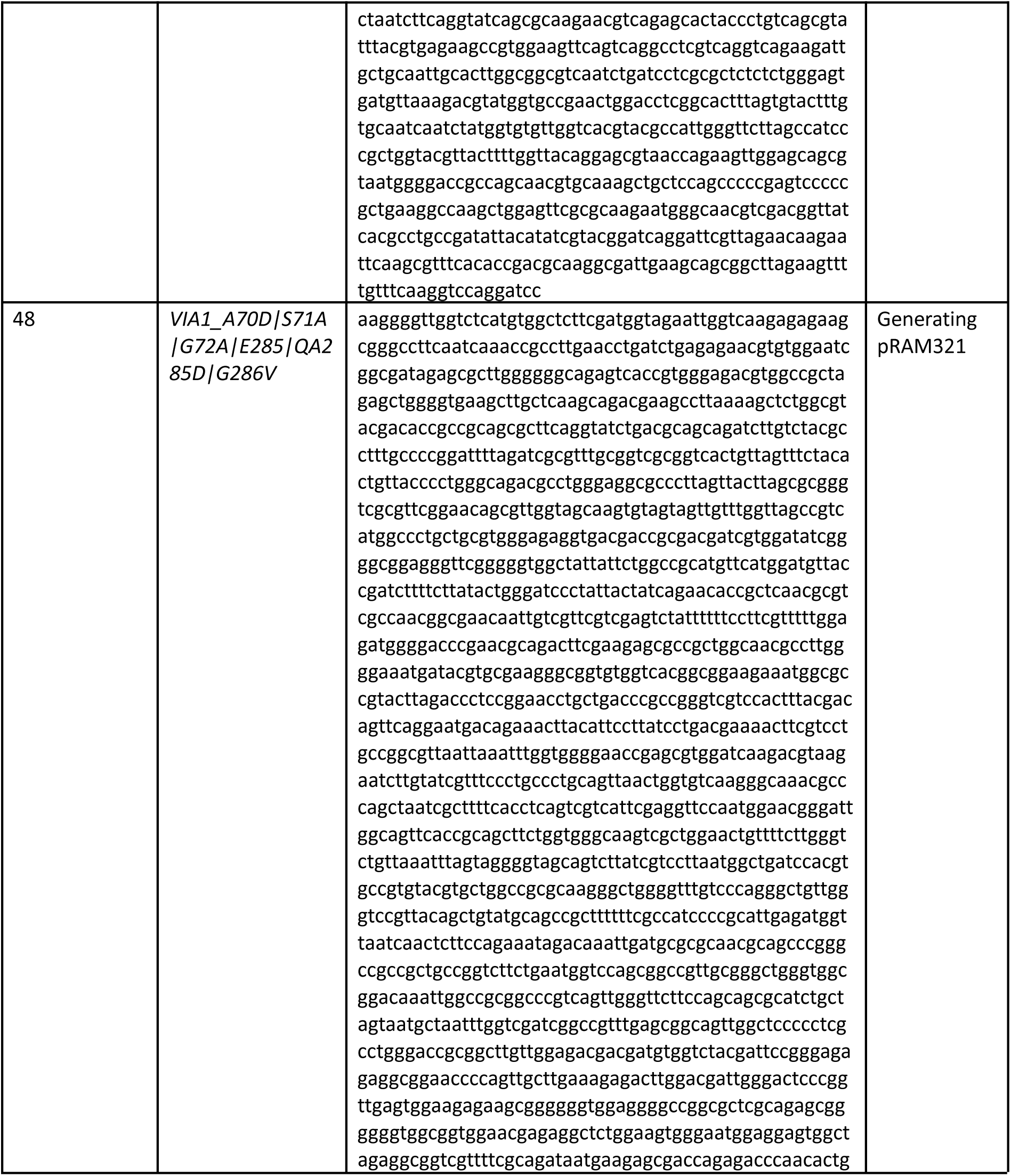
List of Twist fragments/vectors used in this study.

**Table S5.**
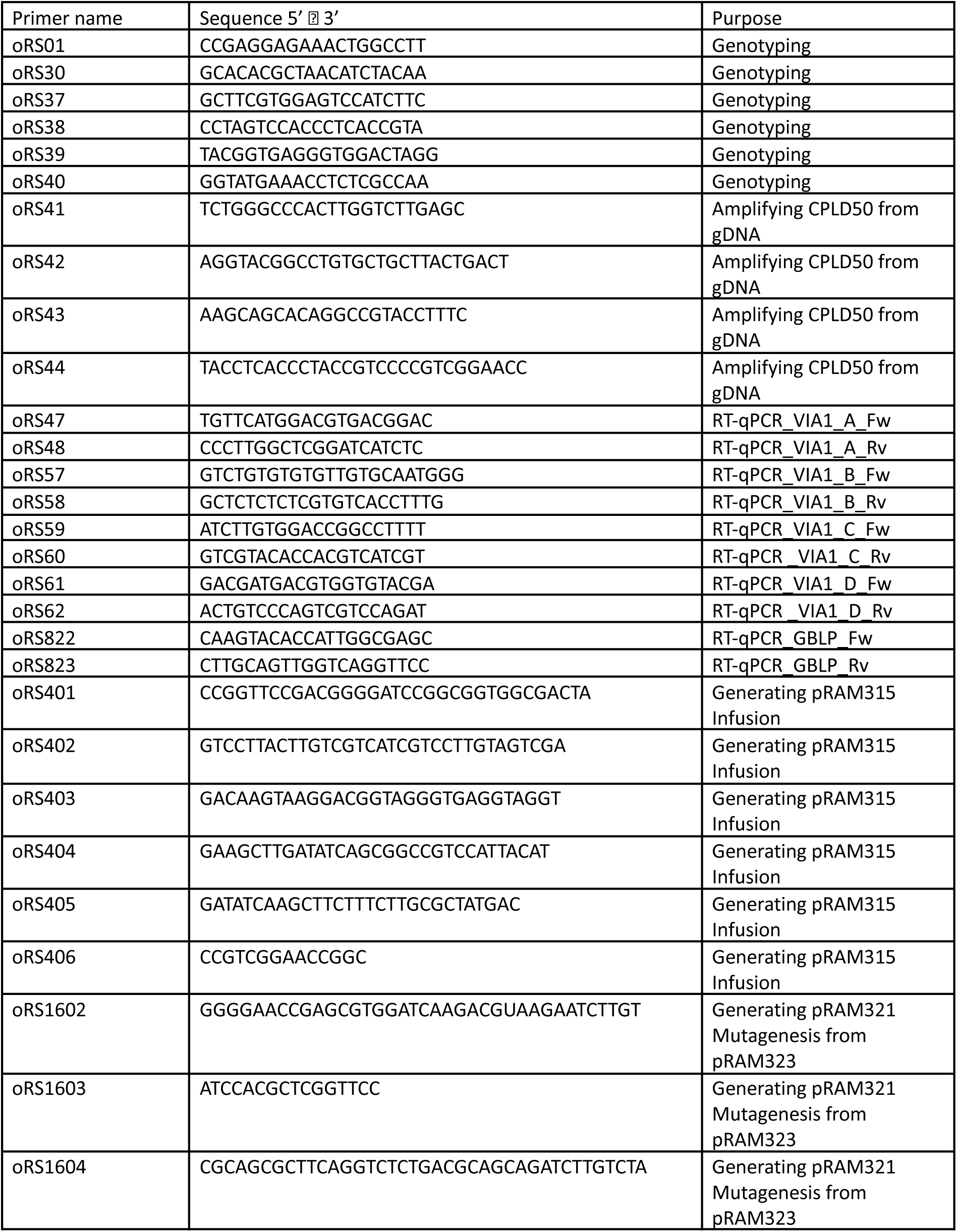
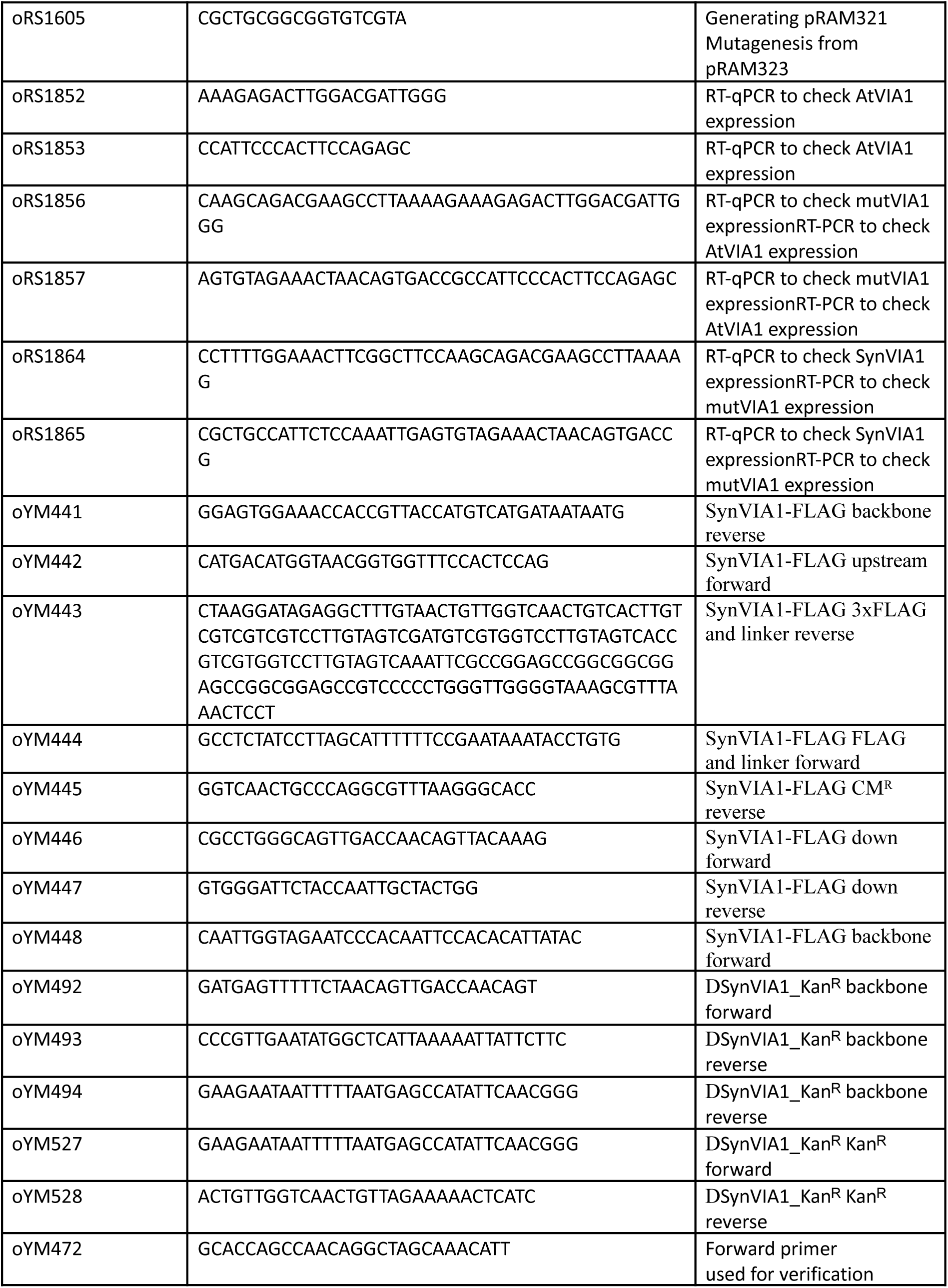
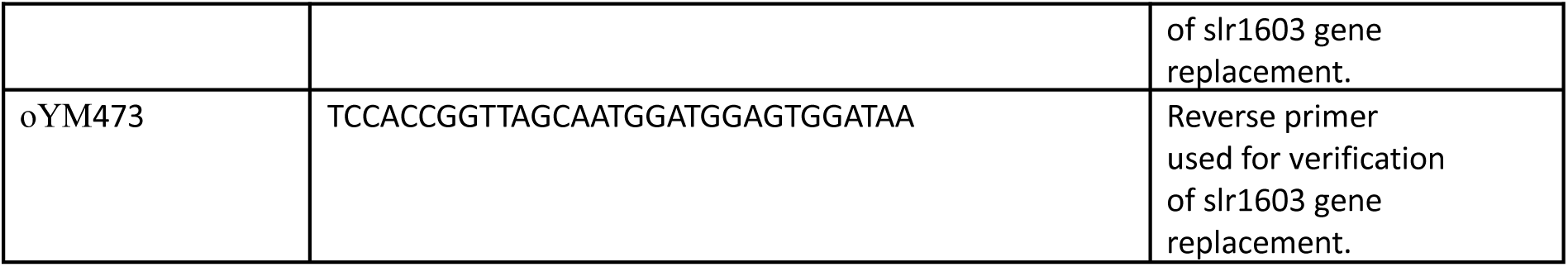
List of oligonucleotides used in this study.

